# Disrupted O-GalNAc glycosylation as a mechanism and biomarker of *SLC35A2*-associated epilepsy

**DOI:** 10.64898/2026.03.02.708854

**Authors:** Robert G. Mealer, James J. Anderson, Sheridan L. Smith, Brianna M. Masters, Samuel H. Barth, Kaitlyn D. J. Huizar, Sahibjot Sran, Hyojung Yoon, Amanda Ringland, Savannah J. Muron, MaryAnn E. Bowyer, Alissa M. D’Gama, Annapurna Poduri, Hart G. W. Lidov, Edward Yang, Julia Furnari, Peter D. Canoll, Adam P. Ostendorf, Daniel C. Koboldt, Christopher R. Pierson, Diana L. Thomas, Benjamin D. Philpot, Maxence Noel, Richard D. Cummings, Erin L. Heinzen, Tracy A. Bedrosian

## Abstract

Rare germline and somatic variants in *SLC35A2* cause a spectrum of severe glycosylation disorders that commonly present with epilepsy. *SLC35A2* encodes the Golgi transporter for UDP-galactose, but how its deficiency leads to severe neurodevelopmental disorders is unknown. Using a mouse model deficient for *Slc35a2* in the forebrain, we identified a specific defect in O-GalNAc glycan synthesis, while other galactose-containing glycoconjugates remained intact. O-GalNAc glycans were absent from their normal location within neuronal tracts of the corpus callosum, and truncated precursors accumulated in the cortex on critical extracellular matrix molecules. Cultured primary neurons lacking *Slc35a2* showed impaired development, hyperexcitability, and impaired O-GalNAc glycosylation. Finally, human brain tissue from cases of *SLC35A2*-associated intractable epilepsy displayed a strong correlation between variant burden and truncated O-GalNAc glycans. These findings provide a mechanistic link between genetic causes of *SLC35A2*-associated epilepsy and protein O-glycosylation that can be targeted for biomarker and therapeutic development.

## Introduction

*SLC35A2* encodes the only known transporter for uridine diphosphate galactose (UDP-Gal), shuttling it from the cytosol to the Golgi lumen. There, UDP-Gal is an essential substrate for all galactosyltransferase enzymes, which covalently transfer galactose to an array of glycoconjugates including glycolipids and glycoproteins to regulate their structure and function. Germline deficiency of *SLC35A2* leads to a severe, X-linked dominant congenital disorder of glycosylation (CDG) termed SLC35A2-CDG, which presents as a developmental and epileptic encephalopathy, though most cases are identified in females, presumably related to prenatal lethality in males which harbor only one copy of the gene^1,2^.

While the expression of *SLC35A2* is low but ubiquitous across nearly all tissues and cell types, the most prevalent and severe phenotypes of SLC35A2-CDG are neurologic, including developmental delay, intellectual disability, and generalized epilepsy^3^. As such, the brain appears uniquely vulnerable to SLC35A2 dysfunction (**Table S1**)^4^. Brain imaging studies of SLC35A2-CDG include cerebral atrophy, delayed myelination, small corpus callosum, and patchy white matter changes reported in many cases^3^. Recently studies have identified somatic (post-zygotic/acquired) variants in *SLC35A2* in male and female individuals with drug-resistant neocortical epilepsy (DRNE), with resected tissue from most of the cases exhibiting a histopathological classification of mild malformation of cortical development with oligodendroglial hyperplasia and epilepsy^5–7^.

Our group recently generated mouse models of *Slc35a2* deficiency using the LoxP system and several Cre driver lines including the forebrain-specific *Emx1* (neurons and glia) and *Olig2* (oligodendrocytes)^8^. *Slc35a2* deletion in *Emx1*-expressing cells led to early lethality, abnormal cortical development, increased oligodendrocyte density, and severe epilepsy that was more pronounced in males than females, mirroring the human SLC35A2-CDG phenotype. Deletion of *Slc35a2* from the *Olig2* lineage resulted in increased oligodendrocyte density and abnormal EEG findings but did not cause a severe epilepsy phenotype or profound lethality. These *Olig2*-specific results were recently replicated by an independent group^9^.

The mechanism by which *SLC35A2* deficiency causes disease, particularly in the nervous system, is presumably through impaired galactosylation. Recent functional studies identified impaired neuronal migration in mice^8,10,11^ and abnormal neuronal development in human induced pluripotent stem cell (hiPSC)-derived neurons with reduced N-glycan formation^12^.

However, the connection between specific glycosylation changes and these neurodevelopmental processes remains unclear. Surprisingly, serum markers of abnormal protein N-glycosylation are mostly absent in SLC35A2-CDGs, or in some cases normalize after infancy^1,2^. Different mammalian cell models of SLC35A2-deficiency have been generated that show isolated glycosylation changes in some pathways but not others^2,13–15^. We are unaware of any comprehensive glycosylation studies in mammalian cell types, specifically neurons and glia, that explain the symptomatology of SLC35A2 disorders.

Protein glycosylation in the adult brain is unique from most other tissues^16^. The majority of asparagine (N)-linked glycoproteins have high mannose structures (∼60%), with the remaining complex structures containing large amounts of fucose and bisected N-acetylglucosamine (GlcNAc). Less than 15% of N-glycans in the adult brain contain galactose (Gal), though studies of younger mouse brains have a higher abundance of Gal-containing N-glycans^17,18^.

Interestingly, this pattern appears to be programmed into the differentiation process of neurons. Both glutamatergic and GABAergic neurons derived from hiPSCs show a similar profile with a lack of complex galactosylated N-glycans once mature, while endothelial cells derived from the same iPSC lines have many more galactosylated glycans^19^.

In contrast, nearly all serine/threonine (O)-linked glycans, the most abundant of which are of the N-acetylgalactosamine (GalNAc), or mucin type, contain galactose linked to GalNAc^16^. We recently showed that O-GalNAc glycans in the mouse brain were enriched in neuronal tracts of white matter and present at nodes of Ranvier^20^, in contrast to N-glycans which appear more diffuse and throughout the gray matter^21^. Recent studies also implicate brain O-GalNAc glycans in schizophrenia risk, mouse models of depression, and again of the brain endothelial glycocalyx^18,22–24^. Glycoproteomic experiments also reveal key differences in the carriers of different glycan classes, with N-linked glycoproteins present on a wide range of plasma membrane proteins, receptors, and extracellular matrix molecules^25^, while O-GalNAc glycans were dramatically enriched on a specific class of cell adhesion molecules termed lecticans^20,26^, which are also canonical carriers of chondroitin sulfate proteoglycans. While O-mannose linked glycans, proteoglycans, and glycolipids also contain galactose, the phenotypes of Mendelian disorders associated with these pathways are generally distinct from the epilepsy and intellectual disability caused by SLC35A2 deficiency, though a few exceptions exist^27^. Based on these observations, we hypothesized that O-GalNAc glycans would be more vulnerable to perturbations of galactose conjugation caused by *SLC35A2*-deficiency.

Here, using *Emx1-Slc35a2* conditional knock-out mice, we report that O-GalNAc-linked glycoproteins are disproportionally altered in comparison to other glycoconjugates, specifically N-linked glycoproteins and proteoglycans. Truncated O-GalNAc glycans were identified on important extracellular matrix molecules using glycoproteomics, and primary neurons from these mice had dense staining for truncated O-GalNAc glycans and replicated the hyperexcitability phenotype seen in the disorder. Finally, human brain tissue samples from two cohorts of individuals with intractable *SLC35A2*-epilepsy showed a strong correlation between truncated O-GalNAc glycans and variant allele frequency (VAF) - which were detected only in an area positive for truncated O-GalNAc glycans - which may represent a novel biomarker and treatment target for *SLC35A2*-related epilepsy.

## Results

*SLC35A2* encodes the only known transporter for UDP-Gal from the cytosol to the Golgi lumen, where it is a substrate for galactosyltransferase enzymes, for example T-Synthase (C1GALT1) and its exclusive chaperone Cosmc (C1GALT1C1) (**Fig. 1A**). To detect the presence of different O-GalNAc glycans, we utilized specific carbohydrate-binding lectins *Vicia villosa* agglutinin (VVA) and peanut agglutinin (PNA) (**Fig. 1B**). VVA binds terminal α-linked-N-acetylgalactosamine (GalNAcα1-Ser/Thr, *Tn-antigen*), while PNA binds the Core 1 disaccharide (Galβ1-3GalNAcα1-Ser/Thr, *T-antigen*) without sialic acid. As most brain O-GalNAc glycans contain sialic acid, treatment with Neuraminidase A (NeuA) increases PNA binding and allows select cleavage with O-glycosidase (OGA), which can release the Core 1 disaccharide.

**Figure 1.**
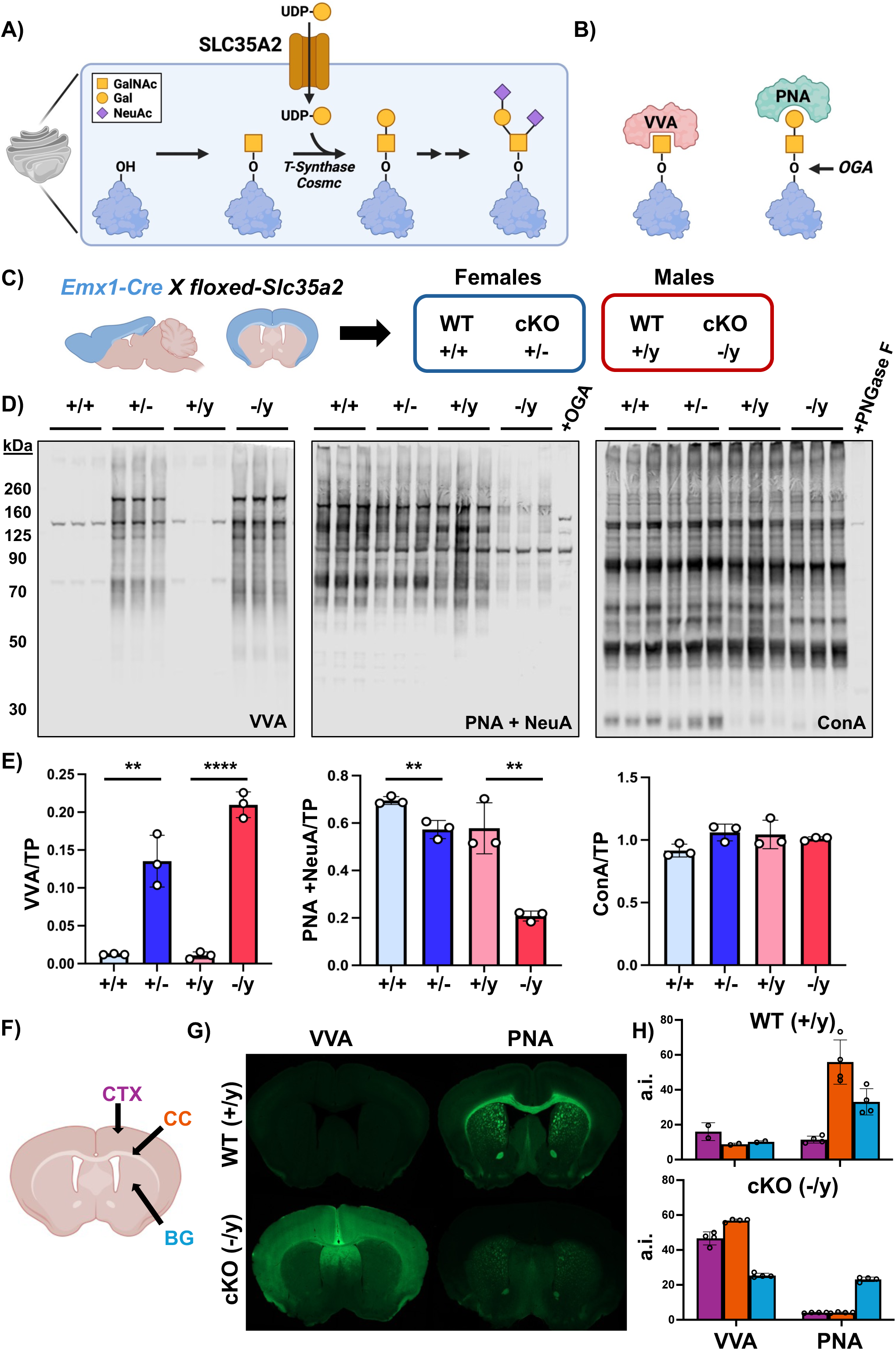
Genetic deletion of *Slc35a2* in the mouse forebrain impairs protein O-GalNAc glycosylation. **A)** *SLC35A2* encodes a transporter for UDP-galactose, which is required for the synthesis of O-GalNAc glycans. **B)** The lectin VVA binds terminal alpha-GalNAc (Tn-antigen), while PNA binds terminal Gal of Core 1 O-GalNAc glycans (T-antigen), which can be removed by O-glycosidase (OGA). **C)** Breeding strategy of *Emx1-Cre* mice with floxed-*Slc35a2* mice, with blue regions highlighting the forebrain expression *Emx1*. **D)** VVA binding is minimal in protein blots of wild-type mouse cortex; female (+/+) and male (+/y) with floxed-*Slc35a2*. In contrast, mice lacking *Slc35a2* in cells expressing the forebrain-specific promoter Emx1 (*Emx1-Cre*) display robust VVA binding in both females (+/-) and males (-/y), consistent with inhibition of O-GalNAc extension. Binding of PNA after Neuraminidase A (NeuA) treatment was significantly reduced in both +/- and -/y lines, and specificity of PNA binding was confirmed via OGA treatment of a +/+ sample. ConA, which binds all N-glycans, was unaffected in all mouse lines, and specificity was confirmed using peptide N-glycosidase F (PNGase F) treatment of a +/+ sample. Each lane contains 15 µg cortical protein lysate from an individual mouse of the corresponding genotype. Experiment was replicated three times with similar results. **E)** Quantification of lectin binding from D normalized to total protein (TP) staining within the same lane of each blot. Data presented as the mean binding intensity +/- SEM, with data points for each individual mouse included, N = 3/genotype. One-way ANOVA confirmed significant group differences (*p* < 0.05) for VVA and PNA but not ConA, followed by post hoc Student’s t-tests to confirm genotype effects in each sex: \*\**p*-value = 0.01, \*\*\*\**p*-value = 0.0001. **F)** Schematic of coronal mouse brain section highlighting the cortex (CTX), corpus callosum (CC), and basal ganglia (BG). **G)** VVA binding is minimal across all regions in wildtype mouse cortex (+/y), while PNA binding localizes to neuronal tracts of the CC and axon bundles in the BG. In contrast, mice lacking forebrain expression of *Slc35a2* (-/y) display a build-up of truncated O-GalNAc glycans in cortex that fail to properly localize. Mature O-GalNAc glycans are entirely absent from the CTX and CC, but preserved in the BG, which does not express *Emx1*-Cre. **H)** Relative quantification of lectin binding across brain regions (CTX, CC, BG) from G. Data presented as the mean binding intensity +/- SEM, with individual points representing the average of 6 measures across independent mice. Colored bars correspond with the specific brain region indicated in (F): CTX, purple; CC, orange; BG, blue. N = 2 for +/y VVA; N = 4 for +/- PNA, -/y VVA, and -/y PNA. Fluorescence was quantified as arbitrary intensity (a.i.) after subtracting background signal from each sample.

### Loss of Slc35a2 in forebrain preferentially impairs protein O-GalNAc glycosylation

To determine the effect of *Slc35a2* deletion on brain glycosylation, we crossed homozygous floxed *Slc35a2* female mice with homozygous male mice expressing Cre recombinase under the forebrain-specific promoter *Emx1* (*Emx1-Cre*+/-) (**Fig. 1C**) as previously described^8^.

Offspring at postnatal day 21-23 were studied prior to onset of significant lethality^8^. As *Slc35a2* is an X-linked gene, this resulted in unaffected wild-type control mice, both female (+/+) and male (+/y), as well as mice lacking *Slc35a2* in cells expressing *Emx1*, both heterozygous females (+/-) and hemizygous knockout males (-/y). Of note, homozygous deleted females cannot be generated due to an apparent lack of fertility of heterozygous deleted females and the early lethality of hemizygous knockout males^8^.

VVA, which does not bind glycoproteins in most normal tissues including brain, was absent in control (+/+, +/y) mice on protein blots. However, both +/- and -/y mice showed robust VVA binding that was observed as discrete bands along with a smear pattern (**Fig. 1D**); these were significantly different from control mice (*p*-value: +/+ vs. +/- = 0.0034; +/y vs. -/y = <0.0001) (**Fig. 1E**). Analysis of PNA binding after treatment with NeuA showed a small but significant decrease in female heterozygous deleted mice (+/-) compared to controls (+/+), while -/y cortex showed a near total loss of PNA binding and a pattern similar to control cortex (+/+) treated with OGA to remove all Core 1 O-glycans (*p*-value: +/+ vs. +/- = 0.0071; +/y vs. -/y = 0.0043). In contrast, binding of Concanavalin A (ConA), which recognizes the mannose-rich core motif in most types of N-glycans, showed no change in binding in mice with *Slc35a2* deletion.

Studies using additional lectins that bind terminal features of N-glycans demonstrated that the *Slc35a2* genotype had no significant effect on binding of *Galanthus nivalis* lectin (GNL; high mannose), *Phaseolus vulgaris* erythroagglutinin (PHA-E; bisected-type N-glycans), and *Ricinus communis* agglutinin (RCA-I; galactosylated N-glycans) (**Fig. S1**). *Aleuria aurantia* lectin (AAL) binds both core α1,6- and the antennary α1,3-linked fucose, while *Lens culinaris* agglutinin (LCA) only binds core α1,6-linked fucose. A small but significant reduction of AAL binding was seen in hemizygous deleted male mice (-/y) compared to controls, while no differences were seen for LCA. This result is consistent with *Slc35a2* deletion affecting the small amount of antennary galactose present on N-glycans, as the FUT9 enzyme which generates α1,3-linked fucose in the brain requires the presence of terminal β1,4-galactose for activity. Similarly, *Griffonia simplicifolia* lectin-II (GSL-II) binding, which detects terminal N-acetylglucosamine (GlcNAc) of N-glycans lacking galactose, was of low intensity but increased in *Slc35a2*-deleted cortex. Finally, changes in PNA binding were only present in the cortex and not the cerebellum, consistent with the forebrain-specific expression of *Emx1* (**Fig. S2**). These findings suggest *Slc35a2* deletion in *Emx1*-expressing cells causes predominantly O-GalNAc changes in mouse forebrain, consistent with the high abundance of galactose on O-GalNAc glycans relative to N-glycans.

### O-GalNAc and chondroitin sulfate modifications are present on the same protein carriers

Other classes of glycoconjugates utilize UDP-Gal during synthesis, such as large chondroitin sulfate proteoglycans (CSPGs) which contain galactose residues in the linker region followed by repeating disaccharide units. As such, we asked whether their synthesis may also be impaired by *Slc35a2*-deletion. The lectin *Wisteria floribunda* agglutinin (WFA) is frequently used to bind the terminal GalNAc of CSPGs in the brain, particularly those associated with perineuronal nets (PNNs)^28,29^. However, lectin microarray studies confirm that WFA can also bind terminal GalNAc of glycoproteins^30^. Surprisingly, we found an overall increase in WFA binding in *Slc35a2* deleted mice, noting that the WFA smear below 260 kDa resembled that of VVA (**Fig. S3**). Quantifying the WFA signal above and below 260 kDa showed that the lower molecular weight proteins were driving the increased signal in *Slc35a2*-deleted mice, whereas the signal from higher molecular weight proteins, i.e., CSPGs, was unchanged. These results indicate that WFA is likely detecting terminal O-GalNAc structures of glycoproteins similar to VVA while proteoglycans remained intact.

To further investigate the potential impact of *Slc35a2* deletion on glycoproteins/proteoglycans, we performed combinational glycosidase digestions followed by protein blotting (**Fig. S4**). Treatment with chondroitinase ABC (ChABC) to deglycosylate CSPGs resulted in a lower migration pattern of WFA-bound proteoglycans, as expected, which appeared similar between genotypes. However, NeuA treatment in combination with ChABC dramatically increased WFA binding in control cortex, particularly in the fraction above 260 kDa, potentially by exposing more GalNAc specifically on proteoglycans as the signal was insensitive to OGA. This same effect was present in *Slc35a2*-deleted cortex, though appeared slightly blunted compared to controls, consistent with only a small decrease in the level of proteoglycans bound by WFA. This result suggests that many large proteins bound by WFA carry CSPG chains but few O-GalNAc glycans.

In control cortex, blotting for Neurocan (Ncan), a known CSPG carrier which is also modified by O-GalNAc glycans^20^, showed major consolidation of bound protein following ChABC treatment near 55-60 kDa. The intensity of Ncan binding was dramatically increased by NeuA, similar to WFA. However, while OGA had no effect on WFA migration, it resulted in a clear shift of Ncan down to 50 kDa after both ChABC and NeuA treatment, consistent with the presence of O-GalNAc glycans and CSPG chains on the same proteins. In *Slc35a2*-deleted cortex, Ncan appeared similar following ChABC treatment but at a slightly lower molecular weight and had less signal increase following NeuA treatment. Ncan also appeared at the lower migration level (∼50 kDa) without any further decrease following OGA, consistent with near total deficiency of O-GalNAc modifications of Ncan in the *Slc35a2*-deleted cortex. Interestingly, VVA binding in *Slc35a2*-deleted cortex and PNA binding after NeuA treatment in control and *Slc35a2*-deleted cortex increased following ChABC treatment. This suggests that the removal of the CSPG chains may expose additional O-GalNAc glycans for lectin binding, and again confirming that CSPGs and O-GalNAc glycans are on the same protein carriers. Total protein loads for each blot confirm both appropriate loading conditions and the presence of each enzyme treatment. Taken together, the results indicate that *Slc35a2* deletion results in loss of galactose on GalNAc-containing O-glycans as the Core 1 disaccharide but does not appreciably affect the galactose requirement for biosynthesis of CSPGs.

### Truncated O-GalNAc glycans accumulate in Slc35a2-deleted cortex and do not properly localize

Based on its specific expression in dorsal forebrain neurons (**Fig. 1C**), we suspected that *Emx1*-Cre deletion of floxed *Slc35a2* would result in a loss of galactosylated glycans in the cerebral cortex and its projections into neuronal tracts such as the corpus callosum, while sparing those in more rostral structures including the basal ganglia (**Fig. 1F**). VVA binding was absent throughout the control brain, but robust and diffuse in the cortex and septal nucleus of *Emx1*-Cre *Slc35a2*-deleted mice (**Fig. 1G, 1H**). PNA binding, which is most robust in the corpus callosum of control mice, was absent in *Slc35a2*-deleted mice, while PNA binding to the neuronal bundles of the basal ganglia appeared unchanged. The binding patterns of multiple N-glycan binding lectins, including ConA, AAL, and RCA, remained relatively unchanged in the *Emx1*-Cre *Slc35a2*-deleted cortex (**Fig. S5**), consistent with our lectin blotting results (**Fig. S1)**. Binding of an antibody for myelin basic protein (MBP) to the corpus callosum of *Emx1*-Cre *Slc35a2*-deleted mice was present, consistent with intact white matter of the corpus callosum (**Fig. S5**). Further, deletion of floxed *Slc35a2* in oligodendrocytes using the *Olig2*-Cre line had no effect on PNA binding and did not result in VVA positivity at this early time-point. These findings suggest that forebrain deletion of *Slc35a2* results in a dramatic and specific loss of normal O-GalNAc glycans in neuronal projections through an intact corpus callosum, and an accumulation of truncated O-GalNAc structures in the cortex.

### Primary neurons from Slc35a2-deleted mice have altered electrical activity and truncated O-GalNAc glycans

To better understand the distribution and relevance of truncated O-GalNAc glycans, we generated cultures of primary cortical neurons from control and *Slc35a2*-deleted mice. After 11 days *in vitro*, cultures from both genotypes showed similar neuronal morphology and strong binding of the neuron-specific marker MAP2 (**Fig. 2A**). While only trace amounts were present in control cultures, VVA binding in *Slc35a2*-deleted neurons was robust and diffuse (*p*-value: +/y vs. -/y < 0.0001) (**Fig. 2A, 2B**), with most but not all neurons appearing VVA+ (**Fig. 2C**). Changes in PNA binding were not significantly different, though there was a large range of binding seen between cultures, and they were not treated with NeuA (**Fig. S5**).

**Figure 2.**
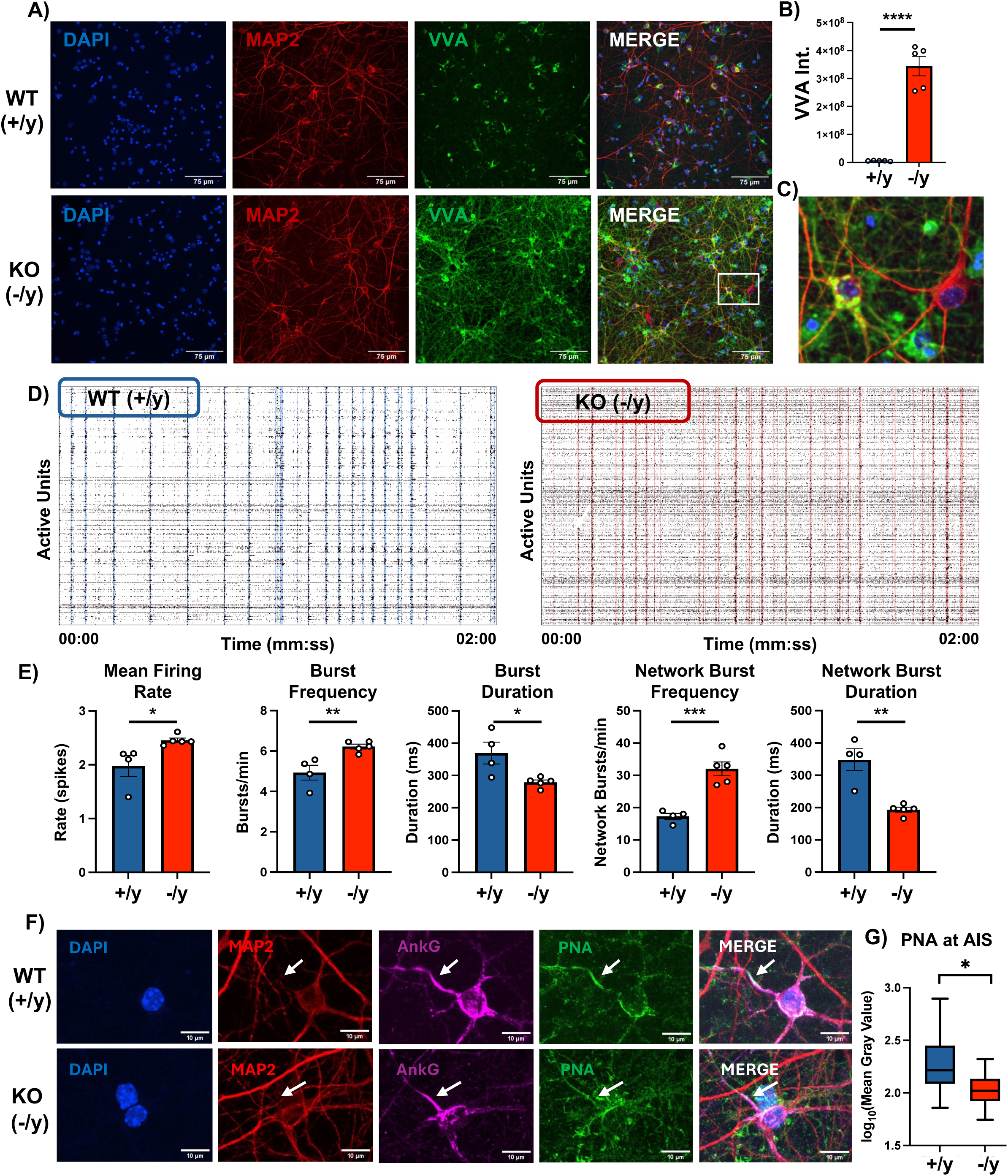
Primary neurons from *Slc35a2*-deleted mice have altered electrical activity and truncated O-GalNAc glycans. **A)** Fluorescent images of +/y and -/y primary neuron cultures at 11 days *in vitro* (DIV11) show robust VVA binding in *Slc35a2*-deleted cultures only. Scale bar = 75 μm. **B)** Quantification of raw integrated density values of VVA fluorescence (VVA Int.). ROIs were selected via automated module. N = 5 independent cultures/genotype. **C)** High magnification of the ROI from A (white box), highlighting that while binding is high in many MAP2+ neurons, some appear VVA negative. **C)** Representative raster plots from 2 minutes of high-density multi-electrode array (HD-MEA) recordings of DIV11 +/y and -/y primary cultures, with active units representing cells firing ≥0.1 spikes/second. Each point represents an individual detected action potential, and colored vertical lines indicate a time when > 20% of active units were recruited during a network bursting event. **E)** Quantification of spontaneous network activity measured by HD-MEA shows that while +/y cultures show clear periods of uniform network activity that recruit many active units, -/y cultures are hyperactive and demonstrate aberrant network activity. N = 4 and 5 independent cultures for +/y and -/y cultures, respectively. **F)** Colocalization of the axon initial segment (AIS) marker Ankyrin G (AnkG) with PNA shows a reduction of properly localized O-GalNAc glycans to the AIS (white arrow, AnkG+/MAP2-). Scale bar = 10μm. **G)** Quantification of the mean PNA value along the AIS from F. N = 3 independent cultures for +/y and -/y cultures, respectively, where 31-32 AISs were measured per condition and average PNA profiles at the AIS were log transformed to fit the normality assumption of the statistical test. Data are shown as mean ± SEM. *p* value = *<0.05, **<0.01, ***<0.001, ****<0.0001 via unpaired two-tailed t-test.

We next assessed the electrophysiologic activity of these primary neurons using multi-electrode arrays. By 11 days *in vitro*, both cultures showed spontaneous electrical activity (**Fig. 2D**). However, comparison of the electrical activity between genotypes showed that *Slc35a2*-deleted neurons had a significantly increased mean firing rate and burst frequency but decreased burst duration (**Fig. 2E, Table S2**). On a network level, increased burst frequency and decreased burst duration were also present, suggesting that *Slc35a2*-deleted neurons display an alteration in network activity, consistent with the observed mouse and human epilepsy phenotypes.

### O-GalNAc enrichment at the axon initial segment is decreased in Slc35a2-deleted cortical neurons

Based on our observation that VVA-bound truncated O-glycans are retained in the cortex of *Slc35a2*-deleted mice (**Fig. 1G**), along with prior findings that mature extended PNA-bound O-glycans are enriched at nodes of Ranvier in neuronal tracts including the corpus callosum^20^, we hypothesized that O-glycans are involved in the formation or function of the axon. The axon initial segment (AIS), a specialized structure involved in initiating action potentials, can be identified by immunofluorescence in cultured neurons as positive for the scaffolding protein ankyrin-G (AnkG), and negative for the microtubule-associated protein 2 (MAP2). Colocalization studies demonstrated an enrichment of PNA binding at the AIS in control neurons, which was significantly reduced in *Slc35a2*-deleted neurons (*p*-value: +/y vs. -/y = 0.049) (**Fig. 2F, 2G**). Thus, while total PNA levels were not significantly reduced by *Slc35a2*-deletion in culture (**Fig. S5**), a deficiency in their specialized localization to the AIS was present at this early developmental stage.

### Lecticans are the primary carriers of truncated O-GalNAc in Slc35a2-deleted cortex

In a prior study, we performed PNA affinity purification after NeuA treatment followed by detailed LC-MS/MS glycoproteomic analysis of a single whole mouse brain^20^. Of the 143 distinct O-GalNAc glycoprotein carriers we found, most were from the lectican family of CSPGs. To determine the carriers of VVA-bound glycoproteins in *Slc35a2*-deleted cortex, we employed a similar approach while incorporating several improvements and controls (**Fig. 3A**). First, biological triplicates of male mice were included as well as a single female heterozygous deleted (+/-) sample. Second, two controls for non-specific binding of glycoproteins in male (-/y) cortex samples were used – an elution control with N-acetylglucosamine (GlcNAc) in place of GalNAc, and a no-lectin control. Third, glycopeptide detection was performed using the ultra-sensitive Thermo Astral LC-MS/MS including a search for specific O-glycan modifications with GalNAc, which are also known as N-acetylhexosamine (HexNAc) as the masses of GalNAc and GlcNAc are identical.

**Figure 3.**
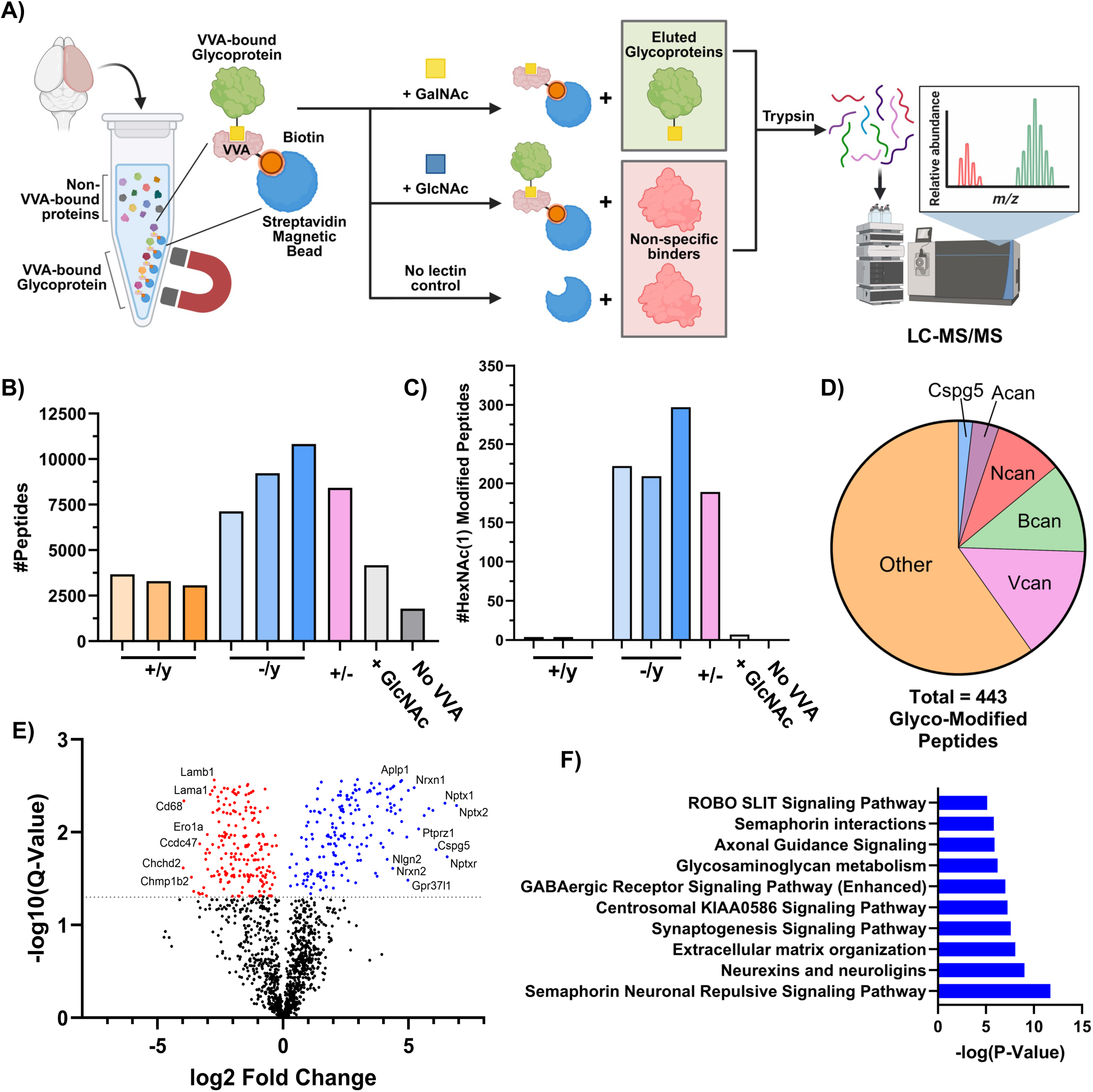
Truncated O-GalNAc in *Slc35a2-*deleted cortex enrich on lecticans and pathways involved in cell adhesion and synapse biology. **A)** Schematic depicting affinity purification and elution of VVA-bound glycoproteins in mouse cortex, including controls for elution (+ GlcNAc) and binding (no lectin), analyzed by LC-MS/MS. **B)** The total number of peptides with spectral count > 0 detected in each sample. Biological triplicates for +/y and -/y were included, as well as a single +/- sample, all eluted with 200 mM GalNAc. Controls for lectin specificity and non-specific binding of a -/y sample were performed in parallel, specifically elution with 200 mM GlcNAc and sample with no VVA lectin. **C)** Among the detected 443 distinct glycopeptides detected, HexNAc(1) modified peptides (+ 203.079 Da) were almost exclusively detected in *Slc35a2-* deleted samples. **D)** HexNAc modified glycopeptides mapped to 60 genes, of which lecticans including Vcan, Bcan, and Ncan were most abundant. **E)** Volcano plot of significantly upregulated (blue) and downregulated (red) proteins in +/y vs -/y cortex, mutants compared to WT. Dotted line indicates significance threshold of *p* = 0.05. **F)** Ingenuity Pathway Analysis (IPA) highlighting the top 10 canonical pathways increased by *Slc35a2* deletion.

Following VVA pulldown of intact glycoproteins, trypsin was used to generate peptide fragments for LC-MS/MS analysis. In total, after excluding contaminants (i.e. albumin, keratins, lectin) 14,753 unique peptides were detected between all the samples (**Fig. 3B**), which mapped to 1,468 proteins (**Fig. S6**). An average of 3,303 peptides were detected in control samples while 8,990 peptides were detected in *Slc35a2*-deleted cortex, with the number of peptides in the elution and lectin controls comparable to control samples. Of these peptides, 464 were found to have a HexNAc modification (**Fig. 3C**), the majority of which were the single HexNAc, though a smaller amount of HexNAc(2) and Hex(1)HexNAc(1) was detected (**Fig. S6**). Of these HexNAc modified peptides, 443 (95%) were found exclusively in the *Slc35a2*-deleted cortices, with very few detected in the three control samples or either control (**Fig. 3C**), confirming the specificity of the pulldown and consistent with lectin blots showing a lack of VVA-bound protein in normal brain tissue (**Fig. 1D**). The 443 HexNAc-modified peptides mapped to 60 proteins, of which over 40% were of the lectican/CSPG family (Ncan, Bcan, Vcan, etc.) (**Fig. 3D**), and this pattern was also observed in the female sample (**Fig. S6**). Gene ontology (GO) analysis of the 60 HexNAc-containing proteins from male mice highlighted many important pathways including extracellular matrix organization and glycosaminoglycan metabolism (**Fig. S6**).

As our purification method precipitated intact glycoproteins in mild detergent and not glycopeptide fragments, the full dataset also included quantitative levels of peptides without HexNAc that may be either (1) on the same protein, (2) a HexNAc-glycoprotein binding partner, or (3) simply non-specific binders. Comparison of peptide levels between genotypes revealed that many proteins had increased and decreased levels in the *Slc35a2*-deleted cortex compared to controls (**Fig. 3E**). As there were essentially no HexNAc-containing proteins in the control samples, the proteins with decreased or unchanged levels are most likely non-specific binders, which is supported by their overall smaller fold change, as well as examples such as Cd68, Lamb1, and Lama1 that are expressed at low levels in the brain. In contrast, the proteins that are increased have a much larger change, with many greater than 16-fold, and include proteins known to have both the truncated HexNAc in this study and extended O-glycans from our previous study^20^, such as Ptprz1, Cspg5, and Nrxn2. Some of the most increased peptides included Nptx1 and Nptx2, which did not have a HexNAc modification. However, their receptor, Nptxr was modified by HexNAc, suggesting that some binding partners were captured by our method. Ingenuity Pathway Analysis (IPA) of the significantly different genes showed that those increased were involved in many relevant pathways including neuronal signaling, binding, ECM, synapse function, and axon guidance, while those decreased were of less clear association with epilepsy phenotypes (**Fig. 3F**).

### O-GalNAc glycans and CSPGs are spatially separated despite both being bound to lecticans

The lectican family of proteoglycans, which includes Ncan, brevican (Bcan), versican (Vcan), and aggrecan (Acan), are some of the most well studied CSPGs in the brain, commonly investigated in the context of PNNs^31,32^. The lectin WFA, which binds terminal GalNAc of CSPGs, is commonly used as a probe to study lecticans and PNNs – however, not all PNNs are WFA positive^28^. Notwithstanding, we used WFA as a probe for CSPGs/PNNs and PNA to bind O-GalNAc glycans to compare their overall spatial distribution in mouse brain and found near total absence of colocalization (**Fig. 4A**). In sagittal sections, PNA binding displayed a similar profile as previously reported^20^, with strong signal in the axon-rich tracts of the corpus callosum, fimbria, and arbor vitae. In contrast, WFA binding was most prominent in nuclear structures including the inferior colliculi, pontine, and deep cerebellar fastigial nuclei, as well as the classical punctate staining of PNNs throughout the cortex. High magnification images of structures with both PNA and WFA signal showed that there was essentially no overlap of PNA and WFA binding. WFA+ cells display a dense ring of binding that excludes nearly all PNA binding (**Fig. 4B**). These results suggest that although CSPGs and O-GalNAc glycans are present on the same protein carriers, they may have distinct roles that impact the localization and function of lecticans or contain different modifications depending on the cell type that expresses the lectican.

**Figure 4.**
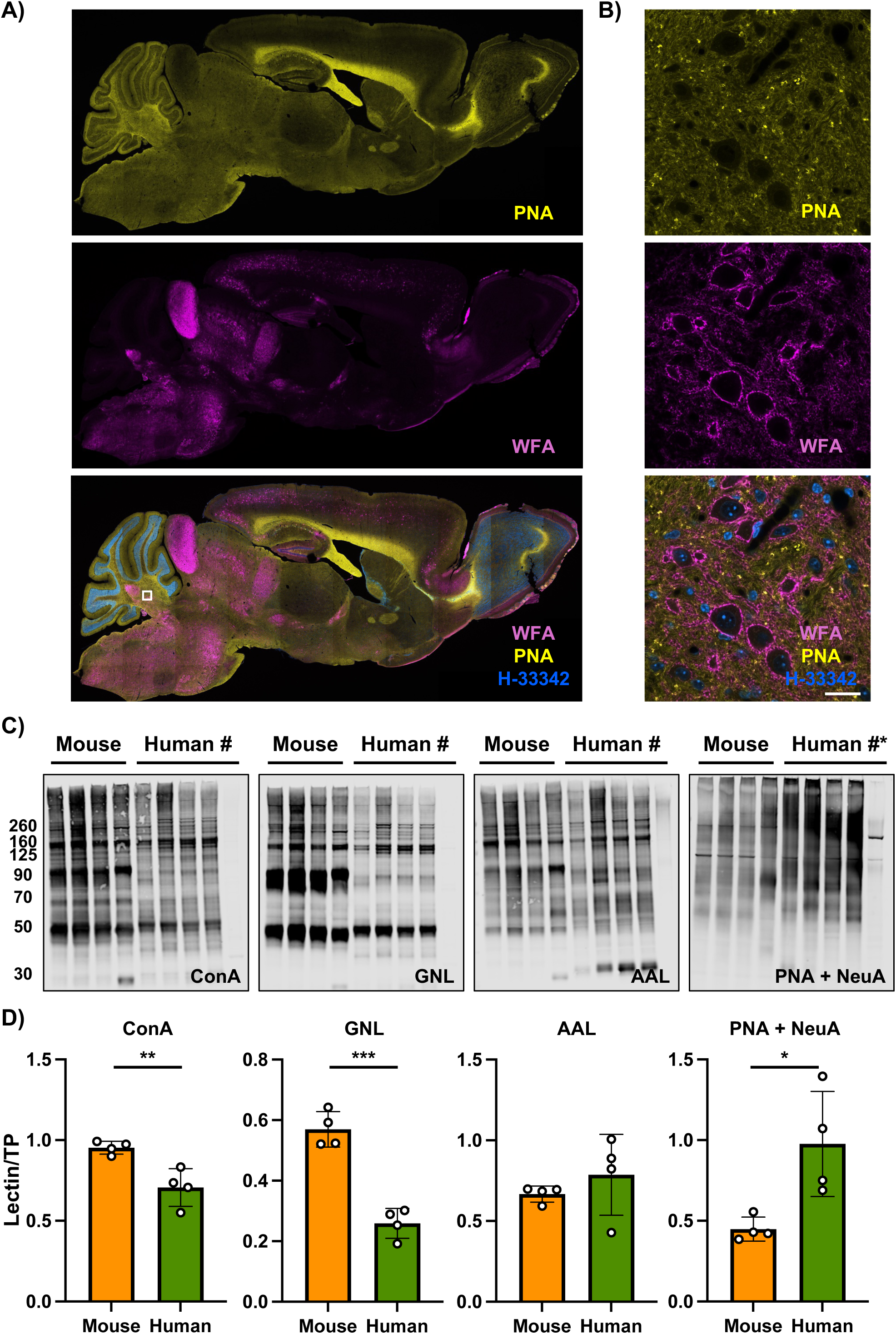
O-GalNAc glycans are spatially separated from CSPGs and more abundant in human cortex than mouse cortex. **A)** Sagittal adult mouse brain sections labeled with WFA for CSPGs and PNA for O-GalNAc glycans display almost no colocalization despite both being bound to lecticans in the brain. **B)** High magnification images of a deep cerebellar nuclei (ROI from A; white box) where both WFA and PNA are present show non-overlapping signals, with WFA densely surrounding cells where PNA is excluded. Scale bar = 20 μm. Nuclei are stained with Hoechst-33324. **C)** ConA, GNL, AAL, and PNA + Neuraminidases A (NeuA) binding (top) from brain cortex samples of wild-type male mice (+/y) and human brain cortex that does not harbor *SLC35A2* mutations. Treatment with PNGase F (#) and O-Glycosidase (#*) of a human sample in the last lane confirms the lectin specificity. **D)** Quantification of lectin binding from C normalized to TP staining within the same lane of each blot. Compared to mouse cortex, human cortex had significantly reduced ConA binding of 27%. GNL staining was decreased even further in humans relative to mice by 55%, while AAL showed no significant difference. In contrast, PNA binding after NeuA treatment was significantly increased in human cortex relative to mice by 44%. Data presented as the mean binding intensity +/- SEM, with data points for each individual mouse and human samples included. N = 4 independent samples per group, with 15 µg cortical protein lysate loaded per lane. Experiment replicated twice with similar results. Student’s t-tests: *p-value = 0.05, **p-value = 0.01, ***p-value = 0.001.

### Human cortex has more O-GalNAc glycans relative to mouse

Changes in the relative abundances of N-glycans between vertebrates have been proposed as possible explanation for the evolutionary divergence of the human brain from other vertebrates, though most LC-MS methods are qualitative or only semi-quantitative. For example, in the dorsolateral prefrontal cortex an increase in the abundance of complex N-glycans from 48% to 52%, and a shift from α2,3-linked sialic acid toward α2,6-linked sialic acid^33^. However, sialylated N-glycans make up less than 1% of all N-glycans in most studies, and we are unable to detect a specific signal in cortex on lectin blot using SNA, which binds α2,6-linked sialic acid^16^. To address quantitative changes of N- and O-glycans between species, we performed lectin blots on wild-type mouse cortex and human cortex from cases of DRNE not caused by variants in *SLC35A2*. Human samples showed a significant decrease in GNL more so than ConA, and no change in AAL (**Fig. 4C, 4D**). These results are consistent with an small increase in complex N-glycans relative to high-mannose and hybrid structures from mice to humans, but do not address the scarcity of galactosylated and sialylated N-glycans as previously reported^16^. Total levels of O-GalNAc glycans bound by PNA after NeuA treatment, as over 90% of O-GalNAc glycans are sialylated, were more than doubled in humans compared to mice (normalized mean - mouse vs human: 0.45 vs 0.98, *p* = 0.02). These results suggest that O-GalNAc glycans may account for a larger amount of the evolutionary divergence of the human brain from other vertebrates compared to N-glycans, and that disruption of UDP-galactose transport by variants in *SLC35A2* may preferentially affect galactose modification of O-glycans over N-glycans.

### Human cortex of SLC35A2-associated drug-resistant neocortical epilepsy shows a diffuse extracellular pattern of truncated O-GalNAc glycans

DRNE may be treated with surgical resection of the seizure focus in some cases, and previous studies of resected tissue have reported somatic variants in *SLC35A2* account for a significant proportion of cases^5,34,35^. We employed confocal imaging and lectin blotting of several such samples and non-*SLC35A2*-associated DRNE controls, as well as performing digital PCR to determine the *SLC35A2* VAF, or what percentage of cells in that piece of tissue contain the variant, of each specimen evaluated (**Fig. 5A, Table S3, S4**). In a case with high VAF (62.6%), we identified strong, but non-uniform, VVA binding across the fixed tissue slice (**Fig. 5B, 5C**). High magnification imaging demonstrated areas of dense VVA staining originating from multiple cells and individual halos around single cells (**Fig. 5D**). To quantify the patchy nature of VVA staining, we measured the relative fluorescence intensity of VVA and DNA in squares ranging from 0.5 mm up to 1.6 mm in diameter, the latter approximating the size of a stereotactic brain biopsy needle (**Fig. 5E, S7**). While the distribution of DNA staining across the brain tissue was relatively consistent, VVA staining had a much larger range, especially in the smallest bin of 0.5 mm. Although autofluorescence of red blood cells in the 488L channel limited our ability to use multiple lectins and colocalization markers in most tissues, the 62.6% VAF sample appeared to have reduced PNA staining in the areas of high VVA binding, suggestive of spatial-restricted inhibition of the O-GalNAc pathway (**Fig. S7**).

**Figure 5.**
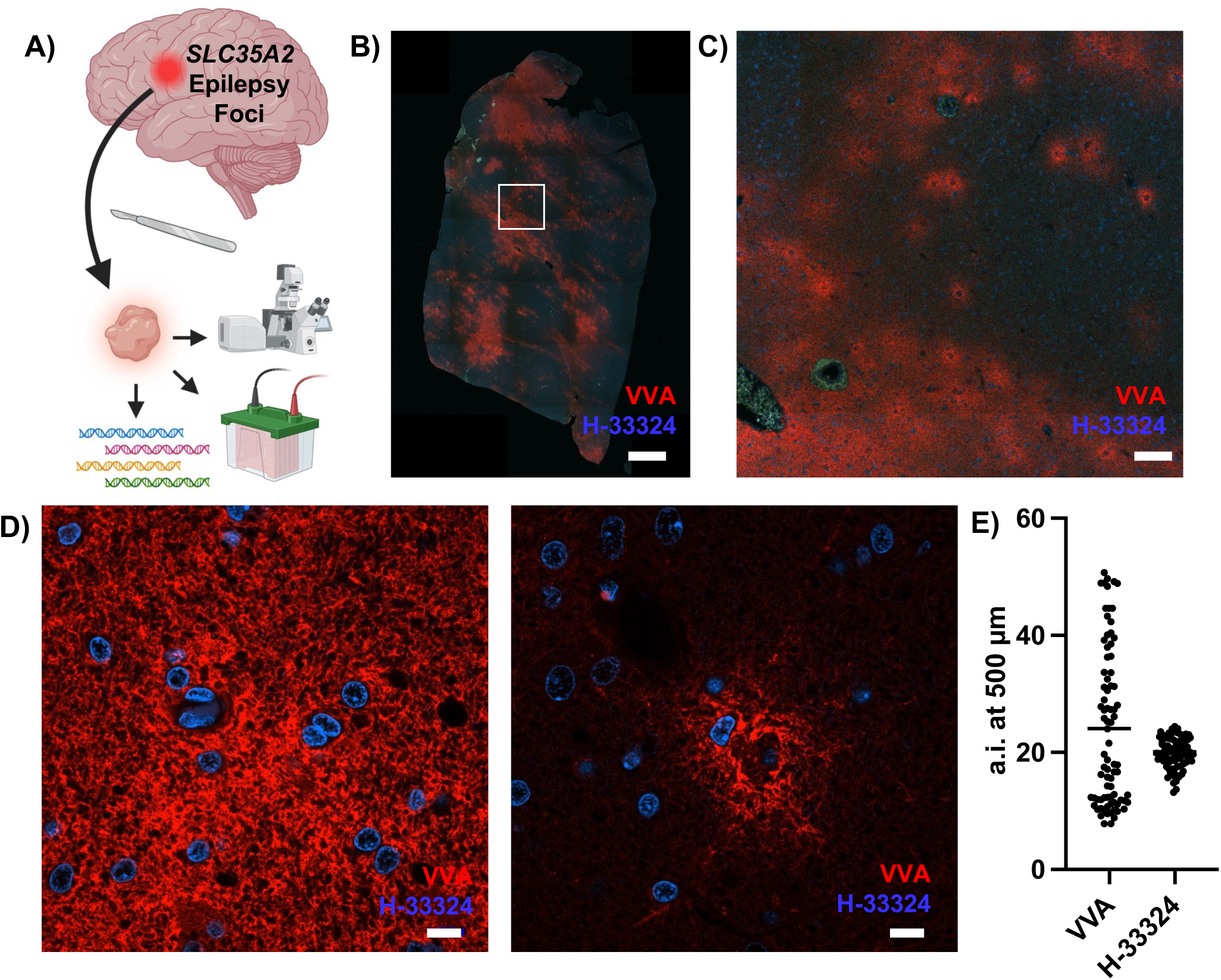
Human brain biopsy tissue from a *SLC35A2*-associated epilepsy case shows diffuse VVA positivity. **A)** Surgically resected tissue from human *SLC35A2*-associated focal epilepsy was analyzed for VVA positivity using fluorescence microscopy, lectin blots, and confirmatory DNA sequencing. **B)** VVA binding across the full tissue block of a high mutation case (VAF = 62.6%) shows variable intensity. Nuclei are stained with Hoechst-33324, and the green signal represents auto-fluorescent red blood cells in the 488ƛ channel. Scale bar = 1 mm. **C)** ROI from B shows scattered binding, with areas of both isolated and diffuse signal. Autofluorescent red blood cells can be seen in the lumen of vessels at the 488ƛ channel. Scale bar = 100 µm. **D)** High magnification images of VVA staining from the same sample, which shows areas of diffuse mesh-like binding around multiple cells and some individual cells. Scale bar = 10 µm. Complementary images with individual channels are available in the Supplementary Material. **E)** Random 500 x 500 µm fluorescent intensity measures of VVA and H-33324 signal from B, plotted as arbitrary intensity (a.i.) normalized to average background signal adjacent to tissue block. Bar indicates median, N = 82 and 80.

To gather additional cytoarchitectural information, we next employed 3,3’-diaminobenzidine (DAB) detection of lectins in fixed H&E-stained tissue in combination with laser capture microdissection (LCM) to simultaneously measure the *SLC35A2*-variant burden (**Fig. 6A**). In a sample with a VAF of 6.3-24.4%, *SLC35A2* amplicon sequencing after LCM from areas that were either VVA- or VVA+ demonstrated that variant sequences were only detectable in VVA+ areas despite equal reference sequence coverage, suggesting that the presence of truncated O-GalNAc glycans are specific for *SLC35A2* dysfunction (**Fig. 6B**). We observed patchy VVA positivity which was spread throughout the white and gray matter (**Fig. 6C**), with similar observations in a second patient sample with a VAF of 2.5-11% (**Fig. S7**). VVA binding during DAB staining was prevented by preincubation of the lectin with the inhibitory sugar GalNAc (200 mM), confirming specificity, and no VVA positivity was detected in a control DRNE sample without *SLC35A2* variants. In addition, the distribution of ConA binding in both samples appeared uniform, consistent with a specific disruption of the O-GalNAc pathway caused by *SLC35A2* variants. High magnification of VVA+ areas revealed binding to both the soma and processes of dysmorphic neurons in low density areas, as well as areas of diffuse VVA+ and white matter hypercellularity (**Fig. 6D**).

**Figure 6.**
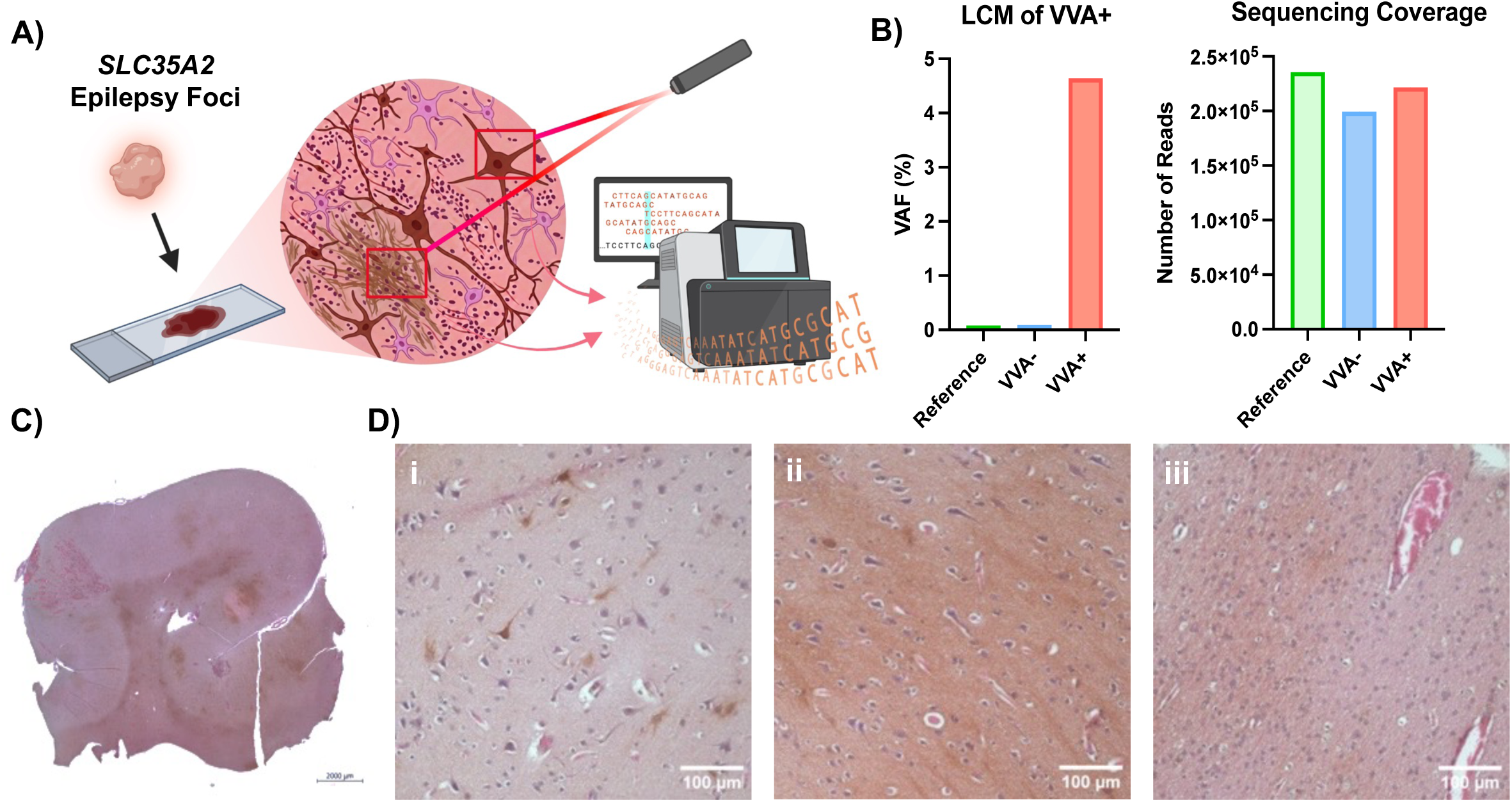
Human brain biopsy tissue from *SLC35A2*-associated epilepsy cases reveals unique histological features and enriches for a patient specific *SLC35A2* variant. **A)** Surgically resected tissue from a *SLC35A2*-associated focal epilepsy patient (VAF = 3.4-5.2%) were analyzed for VVA positivity via lectin-based histochemistry. Using laser capture microdissection (LCM), VVA+ areas were isolated from VVA-areas and both were re-sequenced for the patient specific variant. **B)** Variant allele frequency (VAF) from areas positive and negative for VVA binding was measured using amplicon sequencing of DNA isolated via laser capture microdissection (LCM). Areas of diffuse VVA+ show enrichment for the patient-specific variant in *SLC35A2*, which were absent in both VVA-areas and reference. **C)** Image of the *SLC35A2-*associated epilepsy tissue sample stained with VVA, visualized using DAB histochemistry and counterstained with hematoxylin/eosin. Scale bar = 2000 μm. **D)** High magnification images from (C) of areas of both low density VVA+ with individual cells (i), diffuse high density VVA+ (ii), and hypercellularity in white matter (iii). Scale bar = 100 μm.

### Variant burden in human SLC35A2-associated epilepsy correlates with truncated O-GalNAc glycans but not other glycan classes

To further validate the connection between VAF and truncated O-GalNAc glycans in humans, we performed a blinded analysis of frozen brain tissue from DRNE with and without *SLC35A2* variants via lectin blots (**Table S3**). In a first cohort of 6 *SLC35A2*-cases and 4 controls, VVA+ correctly identified 83% (5/6) of cases after unblinding, with only the lowest VAF sample (2.5%) determined to be VVA-(**Fig. S8**). Normalized VVA signal intensity showed a significant positive correlation with VAF measures from an adjacent but separate tissue block used for lectin blotting (r^2^ = 0.67, *p* = 0.0131) (**Fig. S8**). Given our previous finding of high variability of VVA+ within a single tissue block (**Fig. 5E**), we measured VAF directly using digital PCR from the lysed tissue sample used for lectin blots, which dramatically improved the correlation (r^2^ = 0.98, *p* < 0.0001) (**Fig. S8**). No clear reduction in PNA binding after NeuA treatment was evident, even in the highest VAF samples (**Fig. S8**), and blotting with an additional panel of 6 lectins including GSL-II, which binds glycans with terminal GlcNAc and had a small but significant increase in *Slc35a2*-deleted mouse cortex (**Fig. S1**), showed no correlation between signal intensity and VAF burden in humans (**Fig. S9**). We then replicated the correlation of VVA+ in a separate cohort of *SLC35A2*-associated DRNE again where the VAF was measured directly from the same lysed sample, with the blinded analysis correctly identifying 100% of cases (3/3) and no controls (0/4) (r^2^ = 0.99, *p* < 0.0001) (**Fig. S10**). Finally, we combined all available *SLC35A2*-associated DRNE samples with a VAF ≥ 0.5 (N = 8) and several controls (N = 4) on the same blot (**Fig. 7A**) and confirmed the positive correlation of VVA+ and VAF (r^2^ = 0.98, *p* < 0.0001) (**Fig. 7B**). To address the possibility that the sample with highest VAF (62.6%) was an outlier driving the correlation, analysis excluding this data point remained highly significant (r^2^ = 0.89, *p* < 0.0001) (**Fig. 7C**).

**Figure 7.**
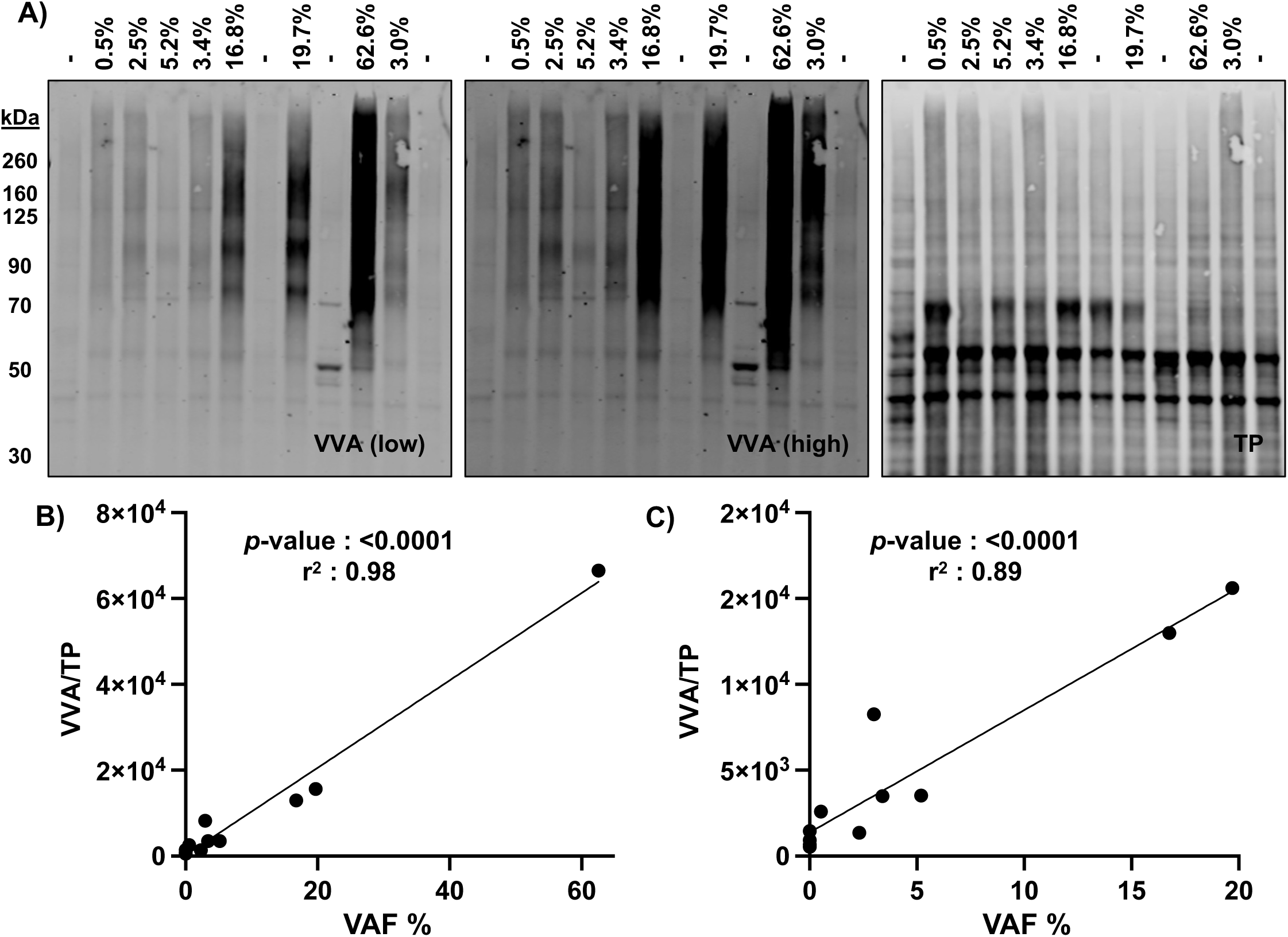
VVA binding in human brain biopsy tissue from two cohorts of *SLC35A2*-associated epilepsy shows a strong linear correlation with variant allele frequency. **A)** VVA lectin blot (both high and low exposure) from human cortex lysate from surgical biopsies of eight cases of *SLC35A2*-associated epilepsy, as well as multiple unrelated controls. The percentage of *SLC35A2* variant allele frequency (VAF) measured on the same same is indicated, as well as controls (-). 15 µg of brain protein lysate was loaded per lane, with total protein (TP) stain indicated in the right panel. **B)** Simple linear regression plot of normalized VVA intensity vs VAF % shows a strong positive linear correlation in *SLC35A2*-associated focal epilepsy. **C)** Linear regression after removal of the highest VAF sample (62.6%) as a potential outlier in B maintains a strong correlation.

## Discussion

In this study, we identify deficient O-GalNAc glycosylation as a critical mechanism in *SLC35A2*-associated disorders using a mouse model, primary mouse neurons, and surgically-resected tissue from patients treated for epilepsy. We hypothesized O-GalNAc glycans would play a major role compared to other galactose-containing glycoconjugates given their unique distribution to the neuronal fiber tracts which are frequently abnormal in MRI studies of SLC35A2-CDG^3^ and the phenotypic overlap of seizures and intellectual disability commonly seen in disorders of protein glycosylation^36,37^. In each experiment, deficient UDP-Gal transport led to an accumulation of truncated O-GalNAc glycans detected by the lectin VVA, which is normally absent in brain tissue.

In our mouse model with forebrain specific deletion of *Slc35a2*, we observed a dramatic reduction in the quantity of extended cortical O-GalNAc glycans and the build-up of truncated species. The levels of other galactose-containing glycoconjugates appeared mostly unchanged, including N-glycans, which contain relatively few galactosylated species compared to O-GalNAc glycans in the brain. Interestingly, while O-GalNAc glycans localize primarily to neuronal tracts in mature animals, the truncated species remain diffusely distributed in the cortex, suggesting that their extension is required for proper localization of their protein carriers to the axon. These results are supported by findings in primary cortical neurons lacking *Slc35a2*, which have a decreased amount of extended O-GalNAc glycans at the AIS. Further, proteomics confirms that lecticans are the primary carriers of both extended and truncated O-GalNAc glycans, suggesting that the maturation of these structures is required for their proper targeting and localization. A recent study of the O-glycoproteome from neuroblastoma cell lines and purified mouse synaptosomes did not identify lecticans as the primary carriers of O-GalNAc glycans^38^, but this is likely due to the lack of mature axonal tracts where O-GalNAc enrich in these samples.

Lecticans are canonical CSPGs and thought to be critical ECM mediators of many processes including neurite outgrowth, plasticity, and the formation of perineuronal nets^28,29,32^. Blotting experiments and glycoproteomics highlight the connection between these modifications on lecticans, but the dramatic differences in their spatial localization remain unexplored.

Gangliosides, which are critical to neuronal membranes, and galactolipids, the major component of myelin, are both glycolipids that contain galactose^39^. Genetic deletion of most single enzymes involved in ganglioside extension changes the composition of glycolipids but result in usually only minor phenotypes including impaired axonal regeneration and late axonal degeneration^27,40^. An exception to this is *ST3GAL5*, a sialyltransferases for gangliosides which is the genetic cause of salt and pepper regression syndrome with infantile-onset epilepsy in humans, though mice lacking the homolog do not have epilepsy^41^. In mice, genetic deletion of UDP-galactose ceramide galactosyltransferase (*Ugt8*), the enzyme responsible for the first synthetic step of all myelin glycolipids, results in normal appearing myelin with glycolipids containing glucosylceramide in place of galactosylceramide. The mice lacking *Ugt8* do not display epilepsy but instead a demyelination phenotype with decreased conduction velocity, tremor, ataxia, and early death with disorganized myelin and elongated nodes of Ranvier^42,43^.

Myelin appears intact in *Emx1-Cre Slc35a2*-deleted mice, and we previously showed that inhibiting extension of O-GalNAc glycans in neurons but not other cell types results in a small but significant shortening of nodes of Ranvier^16^, the opposite of that seen in *Ugt8* deficient mice. Glycolipids are not readily studied by immunohistochemical approaches for their expression and localization, as fixation causes their removal. While we did not analyze glycolipids in this study, it is an important area of future research.

Other studies have reported glycosylation changes associated with deficient UDP-Gal transport, but not in model systems most relevant to human brain disease. For example, *Drosophila melanogaster* lacking the *SLC35A2* ortholog *Ugalt* showed deficient O-GalNAc glycans on muscles and neuromuscular junctions^44^. However, in drosophila, O-GalNAc glycans are present on laminins and dystroglycan, contain glucuronic acid but no sialic acid, and nearly 20% are the ungalactosylated Tn-antigen^45,46^, all starkly different from mammalian brain O-GalNAc glycans^16^. Studies of immortalized mammalian cell lines have reported variable protein glycosylation differences, primarily in N-glycans and one recent study of glycolipids^2,13–15^, but the glycomes of these cell types are very different from neurons^19^. Past studies identified modest increases in truncated N-glycan glycoforms in SLC35A2 deficient patient brain tissue^47,48^; however, the role of O-glycans has been unexplored. Our group previously identified a small reduction in tri- and tetra-antennary galactosylated N-glycans in hiPSC-derived neurons lacking *SLC35A2*, but O-glycans were not analyzed^12^. Brain O-GalNAc glycans were recently reported to be altered by aging and affect blood-brain barrier function, but this study focused on glycans in the endothelial lumen and not the brain parenchyma^23^.

Our results indicate that loss of UDP-Gal transport caused by SLC35A2 variants primarily leads to a reduction of mature O-GalNAc glycans in neurons, leading to seizures and other symptoms of both acquired (somatic) and congenital disorders. Several possible mechanisms may account for such an effect: (1) some galactose-containing molecules may be dispensable in the brain, as pharmacologic inhibition of N-glycan processing prior to the addition of galactose does not affect branching patterns of cultured neurons^49^; (2) other pathways may substitute UDP-Gal with other substrates, as appears to be the case for myelin glycolipids utilizing UDP-Glc^42^; or (3) specific galactosyltransferases may have higher affinity for UDP-Gal, such that during periods of decreased availability certain glycoconjugates are maintained while O-GalNAc structures are lost. Future studies should address how the specific loss of O-GalNAc glycans can lead to seizure.

Identification of the terminal α-linked-N-acetylgalactosamine (t-αGalNAc) epitope in all model systems of SLC35A2 deficiency has clear translational potential. In human DRNE brain resection samples, t-αGalNAc levels revealed by VVA binding were significantly correlated (r^2^ = 0.98) with *SLC35A2* variant allele frequency. This signal could be used as a diagnostic biomarker for transporter dysfunction in cases suspected to be associated with *SLC35A2* variants and quickly employed as a screen prior to or in place of a direct genetic diagnosis.

Further, ultra-high-sensitive glycoproteomic analysis of CSF or serum in cases of DRNE could identify t-αGalNAc specifically on lecticans prior to surgical resection, which could guide therapeutics such as galactose supplementation which is standard treatment for SLC35A2-CDG^4,50^. Other lectins showed no significant association with SLC35A2 dysfunction except for GSL-II, which was not significant in human samples but slightly elevated in mouse samples.

GSL-II binds terminal GlcNAc, which would be increased in the absence of galactose addition to generate the common N-acetyllactosamine (GlcNAc-Gal) disaccharide of N-glycans. However, the signal to noise ratio of GSL-II was far less than VVA, and its lack of translation to human samples make it a less promising target for biomarker development.

While this work demonstrates that brain O-GalNAc glycosylation is altered by deficiency of the UDP-Gal transporter protein in human samples and mouse models, several questions remain. For example, the role of O-GalNAc glycans in the brain at different developmental timepoints is unknown. Further, the relationship between different modifications present on the same protein, such as CSPGs and O-GalNAc on neurocan, but from potentially different cell types or in different subcellular localizations, remains unexplored. Finally, effective treatments to restore proper O-GalNAc glycosylation are needed to address both acquired and congenital neurologic conditions associated with *SLC35A2* variants.

## Methods

### Human Brain Tissue (Cohort 1)

Resected brain tissue was collected from pediatric patients undergoing neurosurgical treatment for drug-resistant epilepsy at Nationwide Children’s Hospital. All patients were enrolled via written consent on an IRB-approved research protocol. Frozen brain tissue and a paired blood sample from each patient underwent exome sequencing and analysis to identify somatic variants, as previously described^51^. All candidate variants were confirmed by amplicon sequencing. Four patients and variants were reported previously, but 3 are new to this study^51^. Formalin fixed paraffin embedded (FFPE) samples were obtained from a subset of cases where possible.

### Human Brain Tissue (Cohort 2)

Resected brain tissue was collected from adult and pediatric patients undergoing neurosurgical treatment for drug-resistant epilepsy at Boston Children’s Hospital, Duke University Medical Center, Lucile Packard Children’s Hospital at Stanford, and Columbia University Irving Medical Center. All patients were enrolled via written consent on an IRB-approved research protocol. Frozen brain tissue from each patient underwent exome sequencing and analysis to identify somatic variants, as previously described^52,53^. FFPE samples were obtained from a subset of cases where possible.

### Experimental animals

Generation, maintenance, and genotyping of forebrain-specific *Slc35a2*-deleted mice were previously described^8^. In brief, floxed homozygous *Slc35a2* female mice were crossed with *Emx1*-Cre and *Olig2-Cre* male mice obtained from Jackson Laboratory (#005628, #025567). All mouse samples in this study were analyzed at postnatal day 21-23, prior to the onset of significant lethality. All experiments were approved and performed in compliance with the animal care guidelines issued by the National Institutes of Health and the institutional animal care and use committees of Nationwide Children’s Hospital (PI: T.A.B.).

### Lectin blots of human and mouse brain

Lectin blots were performed as previously described^16,20^. In brief, at postnatal day 21, mice were euthanized with an overdose of ketamine (300 mg/kg) and xylazine (15 mg/kg) mixture and cortex tissue was dissected from the brain and snap frozen at −80°C. Brain tissue was lysed according to established protocols and quantified using a Pierce BCA assay (Thermo #23227), and 15 mg of protein was loaded per lane in 4-12% SDS-PAGE gels (Invitrogen #NP0323) after streptavidin pre-clearing (NEB #S1420S). Glycosidase control treatments were performed at 37°C overnight following manufacturers protocols (NEB, Neuraminidase A #P0722L, O-Glycosidase #P0733L, PNGase F #P0704L, and α1-3,4 Fucosidase #P0769S; Chondroitinase ABC, Sigma #C2905-5UN).

Following transfer to nitrocellulose and measurement of total protein concentrations using the LiCOR total protein stain (LiCOR # 926-11011), membranes were blocked in TBS-T with 5% BSA for 1 hour, followed by incubation with biotinylated-lectins 1:3,000 dilutions (Vector Labs: VVA #B1235-2, PNA #B1075-5, GNL #B-1245-2, AAL #B-1395-1, GSL-II #B-1215-2, PHA-E #B-1125-2, LCA #B-1045-5, RCA #B-1085-1, WFA #B-1355-2) or 1:15,000 dilution (ConA #B-1005-5) of anti-neurocan antibody (Invitrogen #PA5-79718) at 1:3,000 at 4°C overnight. Membranes were then washed 3 times in TBS-T for 15 minutes, followed by incubation with the streptavidin-800CW (LiCOR # 926-32230) at 1:25,000 concentration for 30 minutes at room temperature, and washed an additional 3 times in TBS-T for 15 minutes. Blots were imaged with the LiCOR Odyssey CLx System and analyzed using LiCOR Image Studio Software (v5.2.5). Lectin signal intensity for each lane was obtained by selecting an entire ROI for each lane and normalizing against the total protein stain for the same area.

### Lectin fluorescence of mouse brain

Lectin fluorescence was performed as previously described, validating target specificity using competitive carbohydrate haptens^20,21^. In brief, at postnatal day 23, mice were deeply anesthetized with an overdose of ketamine (300 mg/kg) and xylazine (15 mg/kg) mixture and transcardially perfused with ice-cold PBS followed by 4% paraformaldehyde. Brains were removed and postfixed in 4% paraformaldehyde for 24 h. After fixation, brains were incubated in 30% sucrose solution and sections were cut on a freezing microtome (Leica #SM2010R) at 30-μm thickness. Sections were blocked, stained, and washed as above for nitrocellulose blots with the following FITC lectins (Vector Labs, VVA #FL-1231-2, PNA #FL-1071-5, ConA #FL-10011-5, AAL #FL-1391-1, and RCA #FL1081-1) at 25 mg/mL, biotinylated WFA (#B-1355-2) at 25 mg/mL, or primary antibody for myelin basic protein (Cell Signaling #E9P7U) at 1:100 overnight at 4°C. Sections were washed with TBS-T 3 times as above, and secondary streptavidin AF568 (Invitrogen #S11226) or anti-mouse AF555 (Jackson Immuno #717-565-150) at 1:1,000 for 1 hour at room temperature followed by 3 additional washes, with the second wash containing 1:1,000 Hoechst-33342 (Thermo #62249). Sections were mounted with Vectashield Vibrance (Vector Labs #H-1700-10) and imaged using a Zeiss LSM 980 confocal microscope in the Neuroscience Microscopy Core at UNC. Quantification of lectin intensity for VVA and PNA staining in cortex, corpus callosum, and basal ganglia was performed using ImageJ2 (v2.16.0/1.54p). Signal intensity from 3 500×500 pixel boxes in each region bilaterally (18 measurements per mouse) were measured and normalized to the average background signal from 5 tissue-adjacent areas (each corner and bottom middle) within the same slide, and an average signal intensity for each region of each mouse was generated.

### Primary Neuronal Cultures

Primary neurons from postnatal day 0 Emx1-Cre-mediated *Slc35a2-*deleted and floxed control male mice were extracted using the Papain Dissociation System (Worthington Biochemical, cat: LK003150) according to the manufacturer’s protocol. Cells were seeded onto coverslips or 6 well HD-MEA chips (CorePlate, 3Brain) coated with 50-100 μg/mL PDL and maintained in Neurobasal-A media (ThermoFisher) supplemented with B27 (ThermoFisher) and L-glutamine (ThermoFisher). Media was changed every 3-4 days. Cultures on HD-MEA chips were maintained in Neurobasal Media until DIV5, at which time the media was changed to BrainPhys (StemCell Technologies) with SM1 supplement (StemCell Technologies).

### Immunocytochemistry and Image Analysis

Cells coated on coverslips were fixed with 4% paraformaldehyde for 30 minutes and rinsed once with phosphate buffered saline. Fixed cells were incubated in blocking solution comprised of 4% bovine serum albumin in tris-buffered saline with 0.1% Tween-20 (TBS-T) for one hour at room temperature. The samples were incubated in TBS-T for one hour with the following antibodies and lectins: Map2 (Synaptic Systems, #188011, 1:500) and biotinylated-VVA (Vector Labs #B-1235-2, 2mg/mL). To confirm specificity of lectin binding, hapten controls were performed by incubating VVA with N-Acetyl-D-Galactosamine (200mM; Cayman Chemicals, #31728) for 30 minutes prior to its use in primary incubation. Primary antibodies and lectins were removed, and cell samples were washed with TBS-T for 15 minutes, three times. The samples were then incubated in the dark with secondary antibodies conjugated to Alexa Fluor 555, 647, and strepdavidin-488 (Invitrogen, A32773; A32795; S32354, 1:500). Finally, the samples were washed with TBS-T for 15 minutes, three times where the second wash contained the nuclear co-stain DAPI (ThermoFisher) and mounted onto slides with mounting media (Invitrogen). Fluorescent images were acquired on an AXR confocal microscope (Nikon) with consistent settings within experiments.

All image analyses were performed using ImageJ (v.2.16.0). Fluorescence channels underwent preprocessing using rolling ball background subtraction (radius =35 pixels for VVA; 50 pixels for all other channels) and mean filtering on lectin fluorescence channels only (radius = 2 pixels) to enhance features for regions of interest (ROI). Channels were z-projected via maximum intensity and Li’s thresholding algorithm (available as an option in ImageJ) was used to mask the fluorescent signal. The Analyze Particles macro was then used to measure the raw integrated density within each ROI.

To quantify average PNA intensity at the axon initial segment (AIS), we used the following protocol adapted from a previous study^54^ with all images blinded until analysis was completed. A line profile was drawn (width = 1 pixel) in ImageJ, starting at the soma and extending through and beyond the AIS, using the fluorescent signal from AnkG to visually define the boundaries.

Mean gray values for both AnkG and PNA fluorescence channels were recorded using the Plot Profile tool. These values were exported, and all AnkG profiles were smoothed using a 15 point (∼2 μm) sliding mean. Both PNA and AnkG profiles were then normalized to a range between 0 (minimum smoothed fluorescence) and 1 (maximum smoothed fluorescence). The proximal start of the AIS was defined as the point where the smoothed and normalized AnkG profile first reached 0.33; the distal end was defined as the point where the profile declined to 0.33 and did not recover for the remainder of the line profile.

### Multielectrode Array Recordings

Extracellular recordings from postnatal cultures on HD-MEA chips were performed using the HyperCam Alpha system (3Brain). Recordings were carried out at 20 kHz from 1,024 electrodes with 60 µm pitch for 2 minutes under stable conditions (35°C, 5% CO2). All analyses were performed using BrainWave5 software (3Brain). Spike detection was performed using the Precise Timing Spike Detection (PTSD) algorithm with a differential threshold set at 10 times the standard deviation and validated with software artificial intelligence. Spike sorting was performed using K-means clustering and principal component analysis. Spike burst detection was performed using a simple Inter-Spike Interval algorithm, with at least 5 spikes firing together in 100 ms interval. Spike network burst detection was performed using a recruitment-based algorithm, with a valid unit threshold of 0.1 spikes/second, recruiting at least 20% of all valid units with a bin size of 50 ms. Active electrodes were defined as channels with a mean firing rate ≥ 0.10 spikes/sec.

### Glycoproteomics

Mouse cortices were lysed and quantified as above for lectin blots. Biological triplicates for +/y and -/y were included, as well as 1 +/- female, as well as 2 additional -/y samples for elution and binding controls as described below (9 total samples). 500 mg of glycoprotein lysates were precleared with 1 mL streptavidin magnetic beads (NEB #S1420S) for 1 hour at room temperature to remove endogenous biotinylated proteins as previously described. Glycoproteins were then incubated with the volume of 1 mL of washed/precipitated streptavidin magnetic beads (NEB #S1420S) and 100 mg biotinylated VVA (Vector Labs #B1235-2) in a total volume of 500 mL lysis buffer overnight at 4°C. Biotinylated VVA was not added to one -/y sample to control for non-specific binding to the magnetic beads (binding control). VVA-bound glycoproteins were precipitated with a magnet and washed/precipitated 3 times for 15 minutes in 1 mL TBS-T. Glycopeptides were eluted from the VVA-biotin-streptavidin magnetic bead complex with 200 mL TBS-T with 200 mM GalNAc (Sigma #A1795). Glycoproteins from one -/y sample were eluted with 200 mL TBS-T with 200mM GlcNAc (Sigma #A8625) to control for non-specific elution from the complex (elution control).

Samples were then submitted for analysis to the UNC Metabolomics and Proteomics Core for liquid chromatography-mass spectrometry (LC-MS/MS) analysis. For each sample, 200 μl of protein from lysate was precipitated by incubation with 6 volumes of ice-cold acetone at -20°C overnight. The following day, samples were centrifuged at 15,000xg for 15 min at 4°C, supernatant was removed, and the pellet was washed with 100 μL of ice-cold acetone. Pellets were air dried at room temperature for 10 min before being resuspended in 100 μL of 1M urea, 50mM ammonium bicarbonate, pH 8, and left on a shaker for 45 minutes to resolubilize.

Samples were then reduced with 5mM DTT at 37°C for 45 min, then alkylated with 15mM iodoacetamide (IAA) for 45 min in the dark, at room temperature. Samples were then digested with 2 mAu LysC for 3 hours at 37 °C followed by an overnight digestion with 4 ug of trypsin at 37°C. The resulting peptides were acidified to 0.5% trifluoroacetic acid (TFA) and desalted using Pierce desalting spin columns using manufacturer recommendations. Eluates were dried via vacuum centrifugation and resuspended in 2% acetonitrile with 0.1% formic acid prior to determining peptide concentration using the Pierce Quantitative Fluorometric Assay.

Samples were analyzed by LC-MS/MS using a Vanquish Neo coupled to an Orbitrap Astral mass spectrometer (Thermo Scientific). A pooled sample was analyzed at the beginning and end of the sequence. Samples were injected onto an IonOpticks Aurora series 3 C18 column (75 μm id × 15 cm, 1.6 μm particle size; IonOpticks) and separated over a 30 min method. The gradient for separation consisted of 2-30% mobile phase B at a 300 nl/min flow rate, where mobile phase A was 0.1% formic acid in water and mobile phase B consisted of 0.1% formic acid in 80% ACN. The Orbitrap Astral was operated in Data Dependent Acquisition (DDA) mode, in which the most intense precursors were selected within a 0.6s cycle for subsequent HCD fragmentation and Astral MS/MS detection. For full MS scans, m/z was set to 375 – 1500, resolution was set to 180,000, maximum injection time was 5 ms, and AGC target was 300%.

MS/MS scans were acquired in the Astral analyzer using an m/z range of 110 – 2000, a maximum injection time of 2.5 ms, and an AGC target of 100%. The normalized collision energy was set to 30% for HCD, with an isolation window of 2 m/z. Precursors with unknown charge or a charge state of 1 and ≥ 7 were excluded.

Raw data files were processed with Fragpipe (v 22.0) using MSFragger (v 4.1) and IonQuant (1.10.27). The files were searched against the mouse SwissProt database downloaded from Uniprot (containing 17,230 entries, downloaded January 2025), the CRAPome contaminants database, and the VAA-Lectin B4 protein sequence. Within Fragpipe the LFQ-MBR workflow was selected, enzyme specificity was set to “stricttrypsin”, up to two missed cleavages were allowed, the minimum peptide length was set to 7, cysteine carbamidomethylation was set as a fixed modification, methionine oxidation, N-terminal acetylation, HexNAc (+203.079 Da), HexNac(2) (+406.158 Da), and Hex(1)HexNAc(1)(+365.132 Da) on Serine and Threonine residues were set as variable modifications. Both precursor and fragment mass tolerances in MSFragger were set to +/- 20ppm. A false discovery rate (FDR) of 1% was used to filter all data. Downstream analysis was performed in Perseus, including missing value imputation, log2 fold change calculation, Student’s t-tests, and Benjamini-Hochberg correction for multiple hypothesis testing. Proteins identified but not quantified in any sample were removed, and single hits where only one peptide was identified were retained. Log2 fold change, p-value, and q-value calculations were computed before and after imputation. Prior to imputation, data were filtered for at least 2 valid values in either of the control or variant conditions. Ingenuity Pathway Analysis (Qiagen) was used for secondary data analysis including statistical testing and Gene Ontology (GO) analysis. Raw glycoproteomics datasets are available in the ProteomeXchange Consortium via the PRIDE partner repository with the dataset identifier PXD073823.

### DAB Lectin Staining of Human Brain Tissue

FFPE tissue was obtained from two individuals with pathogenic *SLC35A2* variants and one control individual. Slide-mounted, FFPE human brain tissue sections (10 μm) were deparaffinized and then antigen retrieval was performed by incubating slides in citrate buffer (Sigma, #C9999) for 3 minutes at 100°C which was followed by three washes, five minutes each, in TBS-T (0.05% Tween 20). To block endogenous peroxidases, tissue sections were incubated with BLOXALL Endogenous Blocking Solution (VectorLabs, #SP-6000-100) for 10 minutes at room temperature and then washed with TBS for five minutes. Then, samples were incubated with Carbo-Free Blocking Solution (VectorLabs, #SP-5040-125), diluted to working concentration in TBS-T, for 1 hour. Excess serum was removed and endogenous biotin was blocked using a streptavidin/biotin blocking kit (VectorLabs, #SP-2002). Biotinylated lectins ConA (VectorLabs,#B-1005-5, 25 μg/mL) and VVA (VectorLabs, #B-1235-2, 25 μg/mL) were diluted in TBS-T and incubated for 1 hour at room temperature. To confirm specificity of VVA lectin binding, hapten controls were performed by incubating working concentration of VVA with a high concentration of N-Acetyl-D-Galactosamine (200mM; Cayman Chemicals, #31728) for 30 minutes prior to its use. After, the slides were washed with TBS-T three times for five minutes each and then incubated in R.T.U. Vectastain Elite ABC Reagent (VectorLabs, #PK-7100) for 30 minutes at room temperature then washed with TBS for 5 minutes. Lectin binding was visualized by incubating sections for two minutes in complete ImmPACT DAB reagent (VectorLabs, #SK-4105) and immediately rinsing for 5 minutes with two changes of tap water. Slides were counterstained with Hematoxilin and Eosin staining kit (Abcam, #ab245880) following the manufacturers recommendation. Sections were dehydrated with ethanol and coverslipped using VectaMount Express Mounting Medium (VectorLabs, #H-5700-60). Images were acquired on an Axio Imager 2 (Zeiss) using consistent settings within experiments.

### Laser Capture Microdissection

FFPE tissue sections (Patient 52) were stained with VVA and mounted on membrane slides (MMI, #50102). Large areas of diffuse VVA positivity and areas lacking VVA were microdissected using a mmi CellCut System (MMI) and collected in mmi IsolationCaps (MMI, #50204, 0.5mL transparent) for DNA extraction and sequencing.

### Amplicon Sequencing and Analysis

DNA was isolated from the collected samples using a QIAmp DNA Micro Kit (Qiagen, #56304) following the manufacturer’s guide for isolating genomic DNA from laser-microdissected tissues. The isolated genomic DNA (∼30 ng) and human reference DNA GM24143 as a negative control were used as templates for PCR that was performed using Q5 Master Mix (New England BioLabs) and SLC35A2 primers (IDT, 200nM): forward (5’-GGTGAGACCTTTGAGCACTTC-3’) and reverse (5’-AGGGTCCTGGGTGAGAAAGAAA-3’). The reaction was performed under the following conditions: 30 seconds at 98°C, 40 cycles of 10 seconds at 98°C, 20 seconds at 58°C, 20 seconds at 72°C, and final extension of 5 minutes at 72°C. Amplified products were purified using SPRIselect beads (Beckman Coulter, #B23317, 1.8x), followed by end-repair and dA-tailing with NEBNext Ultra II DNA library Prep kit reagents (New England Biolabs, #E7645). This was followed by unique-molecular identifier (UMI) indexed adaptor ligation (IDT). Adaptor ligated samples were purified with two rounds of SPRIselect bead-based cleans (1.0x, 0.9x) and used for library amplification with Q5 Master Mix and Illumina P5/P7 Primer mix. After PCR, the final libraries were purified using SPRIselect beads (0.7x) and libraries were pooled and sequenced on NovaSeq6000. SAMtools mpileup and mpileup2cns VarScan (version 2.3.4) commands were used to generate read counts for variant of interest from raw, de-duplicated BAM files.

### Digital PCR

DNA was isolated from the same cell pellets that were used for the protein isolation to assess VVA binding using GenFind V3 (Beckman Coulter) DNA extraction kit. Variant allele frequencies were assessed in specimens evaluated for the extent of VVA binding using digital PCR. Digital PCR was performed using custom-designed Taqman assays targeting the variant and reference alleles for single nucleotide variants in *SLC35A2*. Both the QuantStudio 3D Digital Real-Time PCR system (Thermo Fisher Scientific) and the QuantStudio Absolute Q Digital PCR System (Thermo Fisher Scientific) were used to quantitatively assess allele fraction per the manufacturer’s protocols.

### Statistical Analysis

Unless otherwise specified, statistical analyses were performed in GraphPad Prism (v10.6.1). For groups of two, unpaired Student’s t-tests were performed assuming equal variance. For groups of 3 or more, a one-way ANOVA was performed, followed by unpaired Student’s t-tests to compare individual group differences. Correlation of VAF % and VVA positivity was performed using simple linear regression. A *p*-value <0.05 was considered statistically significant.

## Acknowledgements/Funding

NIH-NIMH (K08MH128712) to RGM. NIH-NINDS (R01NS129784) to TAB. NIH-NINDS (R01NS094596, R01NS115017, R01NS114122) to ELH. NIH-NINDS (R01NS129914) to BDP. NIH-NINDS (K23NS140397) to AMD. SHB is supported in part by NICHD T32 (5T32HD040127). BMM is supported in part by NINDS T32 (5T32NS007431). SJM is supported in in part by NIGMS T32 (T32GM135128). We would like to thank the UNC Neuroscience Microscopy Core (RRID:SCR_019060), which is supported in part by funding from the NIH-NINDS Neuroscience Center Support Grant P30 NS045892 and the NIH-NICHD Intellectual and Developmental Disabilities Research Center Support Grant P50 HD103573. We would like to thank Laura Herring, Natalie Barker, Thomas Webb, Angie Mordant, Aurora Cabrera, and Scott Lyons for their help with the proteomics work performed in this manuscript. This portion of the research is based in part upon work conducted using the UNC Metabolomics and Proteomics Core Facility, which is supported in part by NCI Center Core Support Grant (2P30CA016086-45) to the UNC Lineberger Comprehensive Cancer Center. We would also like to thank the Boston Children’s Hospital Translational Neuroscience Center Human Neuron Core Repository for Neurological Disorders for their support, and Brenda E. Porter at Stanford University for providing tissue samples from individuals with SLC35A2 epilepsy.

## Author Contributions

RGM - conceptualized the project, performed and supervised glycosylation experiments, and wrote the manuscript; JJA - performed primary neuron culture, MEA, confocal imaging human histology, and LCM; SLS – western blots, data analysis, BMM – glycoproteomic analysis, confocal imaging, data analysis; SHB – confocal imaging, proteoglycan analysis, data analysis; KDJH – glycoprotein isolation for proteomics, SS – confocal imaging and analysis; HY and AR - mouse husbandry and tissue collection; SJM and MEB – digital PCR and data analysis; AMD, AP, HGWL, EY, JF, PDC, BEP, and APO - patients samples and clinical data; DCK – variant analysis; CRP and DLT - neuropathology review of lectin stained tissues; BDP – supervision of proteoglycan analysis, MN and RDC – data analysis and contribution of new reagents/analytical tools; ELH – supervision of digital PCR and human sample analysis, wrote the manuscript; TAB - conceptualized the project, supervised animal experiments, neuronal cultures, MEA, LCM, and human sample analysis, and wrote the manuscript. All authors read and approved of the final manuscript.

## Notes

### Competing Interest Statement

The authors have declared no competing interest.

