## Supplemental Figures for "Disrupted O-GalNAc glycosylation as a mechanism and biomarker of *SLC35A2*-associated epilepsy"

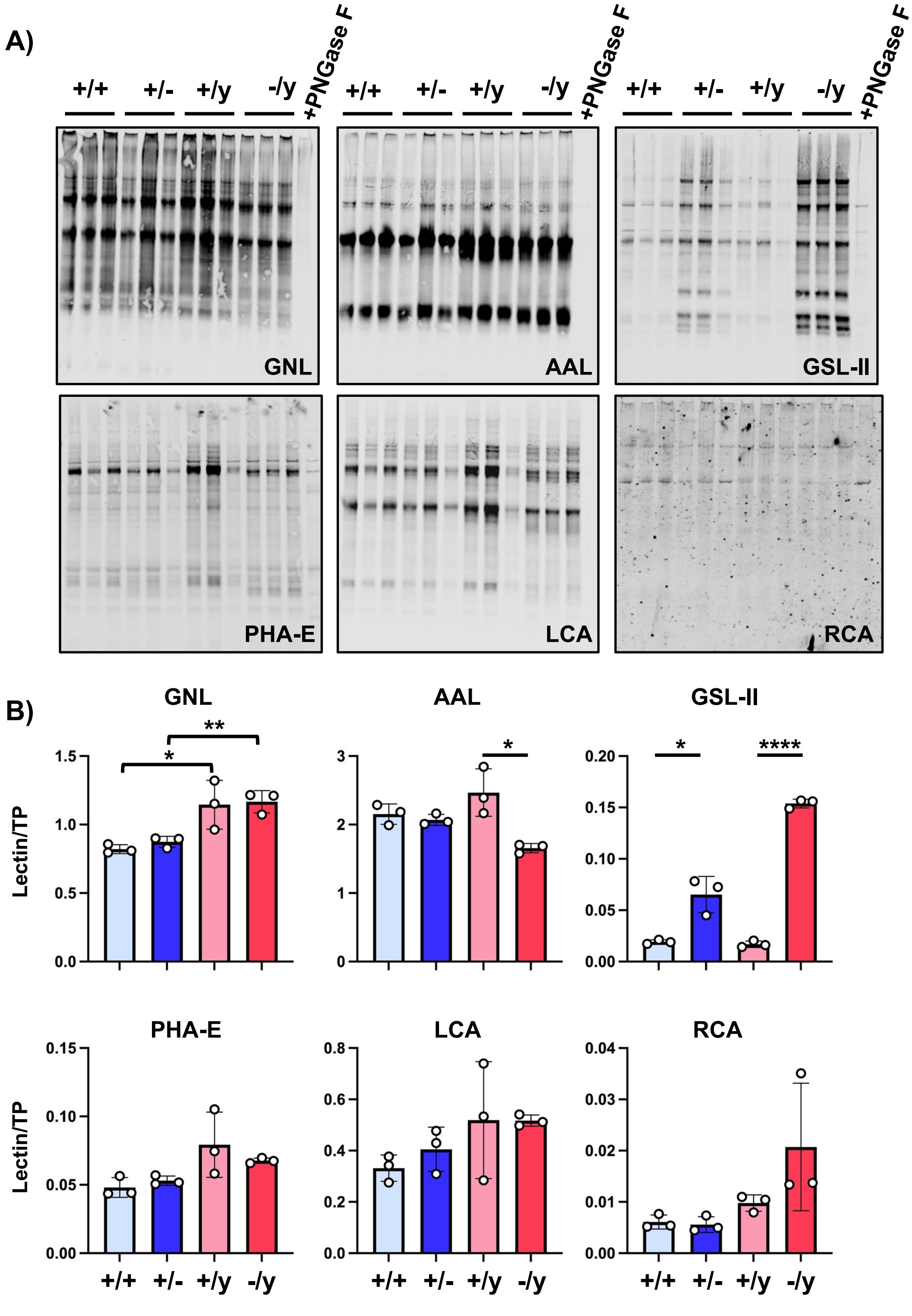


**Supplemental Figure 1. Deletion of *Slc35a2* in *Emx1*-expressing cells has subtle effects on total protein N-glycosylation in cortex. A)** Binding of the lectins GNL, AAL, GSL-II, PHA-E, LCA, and RCA, which are specific for N-glycans, in wild type (+/+, +/y) and *Slc35a2*-deleted (+/-, -/y) mouse cortex. Specificity was confirmed using peptide N-glycosidase F (PNGaseF) treatment of a +/+ sample in the last lane. Each lane contains 15 µg cortical protein lysate from an individual mouse of the corresponding genotype. **B)** Quantification of lectin binding from A normalized to total protein (TP) staining within the same lane of each blot. Data presented as the mean binding intensity +/- SEM, with data points for each individual mouse included. One-way ANOVA confirmed significant group differences ($p < 0.05$) for GNL, AAL and GSL-II. Student's t-tests showed a genotype effect only in males for AAL and both sexes in GSL-II, while group differences in GNL were driven by sex not genotype: **p*-value = 0.05, ***p*-value = 0.01, *****p*-value = 0.0001.

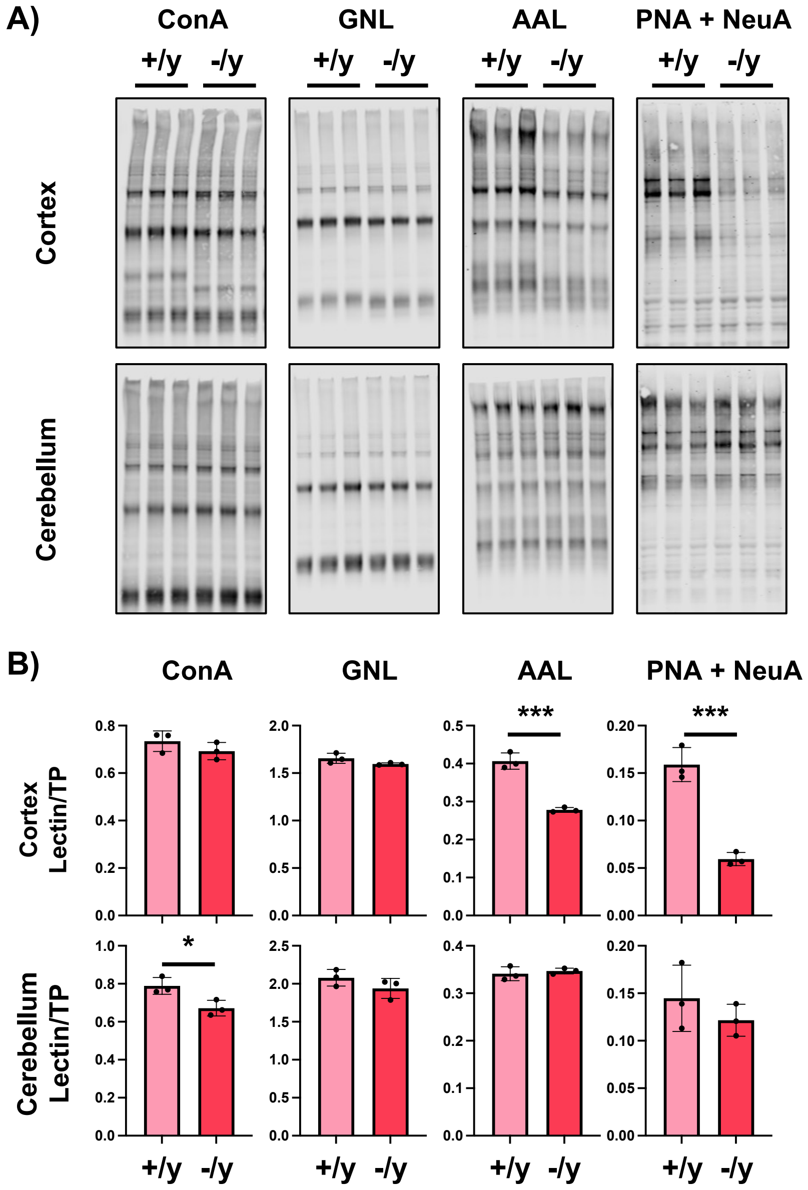


**Supplemental Figure 2. Deletion of *Slc35a2* in *Emx1*-expressing cells inhibits protein O-glycosylation in the cortex but not the cerebellum. A)** ConA, GNL, AAL, and PNA binding in the cortex and cerebellum of wild type (+/y) and *Slc35a2*-deleted (-/y) male mouse cortex. PNA binding was most significantly affected in cortex of *Slc35a2*-deleted mice, but unchanged in the cerebellum. AAL in cortex was significantly reduced, presumably due to the lack of antennary fucose which requires the presence of galactose, as was ConA in cerebellum, though this finding was not significant after correcting for multiple tests. Each lane contains 15 µg cortical protein lysate from an individual mouse of the corresponding genotype. B**)** Quantification of lectin binding from B normalized to total protein (TP) staining within the same lane of each blot. Data presented as the mean binding intensity +/- SEM, with data points for each individual mouse included. Student's t-tests: *p-value = 0.05, **p-value = 0.01, ***p-value = 0.001.

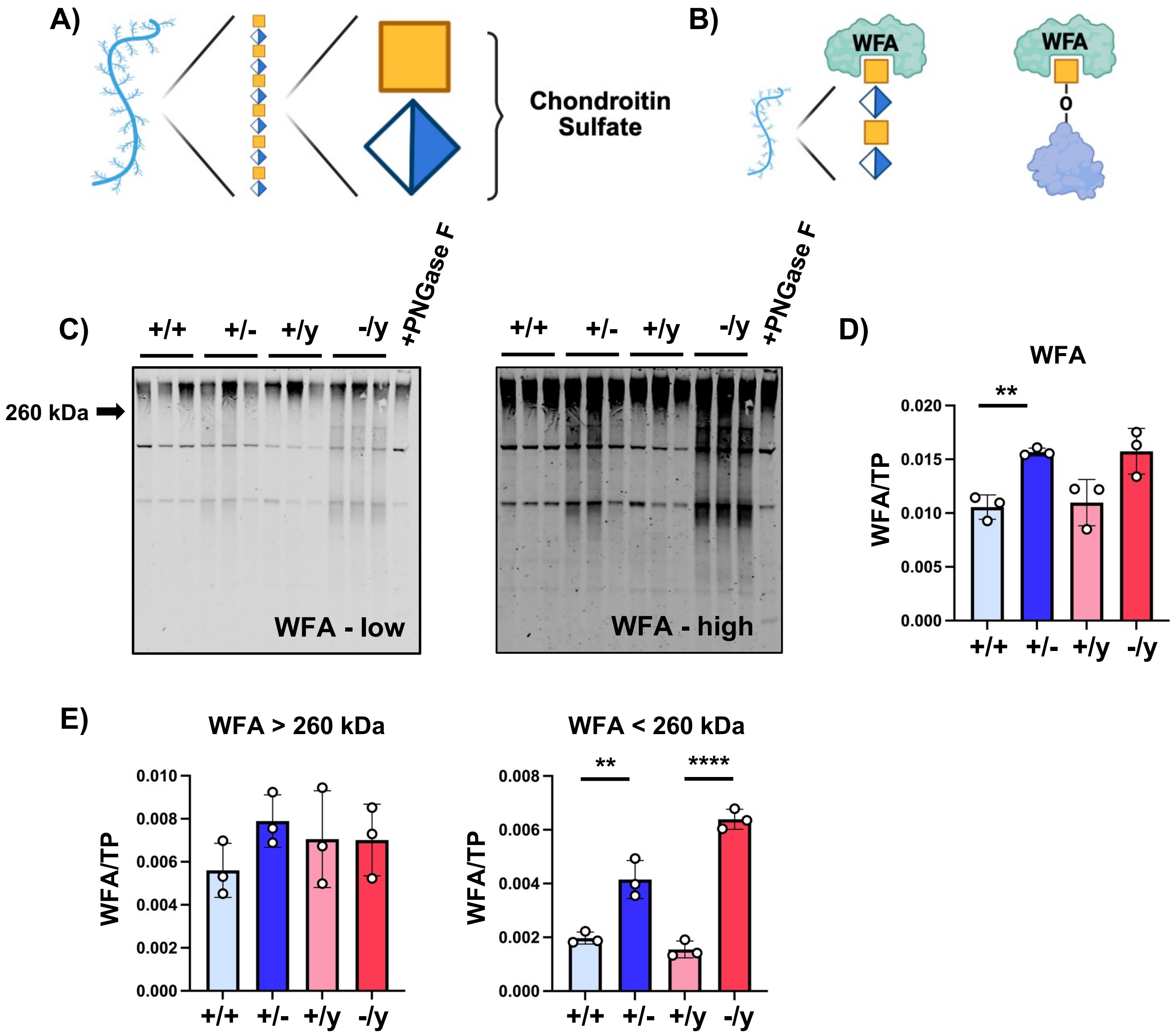


**Supplemental Figure 3. Deletion of *Slc35a2* in *Emx1*-expressing cells increases WFA binding to proteins <260 kDa. A)** Schematic of a typical chondroitin sulfate proteoglycan (CSPG), such as Ncan, which contains repeating disaccharide units of GalNAc and GlcA known as chondroitin sulfate (CS). **B)** The lectin WFA binds terminal alpha-GalNAc, which are commonly found on CSPGs, but is also capable of binding the Tn-antigen. **C)** Binding of WFA to wild-type (+/+, +/y) and *Slc35a2*-deleted (+/-, -/y) mouse cortex, which is insensitive to PNGase F treatment of a +/+ sample in the last lane. Each lane contains 15 µg cortical protein lysate from an individual mouse of the corresponding genotype. A low and high exposure of the same blot is shown. **D)** Quantification of lectin binding from C normalized to total protein (TP) staining within the same lane of each blot. One-way ANOVA confirmed significant group differences ($p < 0.05$), and post hoc Student's t-tests showed a genotype effect only in females, though a similar trend was seen in males. **E)** Quantification of lectin binding from C, separating the WFA signal into proteins above (left panel) or below (right panel) 260 kDa. One-way ANOVA confirmed significant group differences ($p < 0.05$) only for the WFA signal below 260 kDa, and post hoc Student's t-tests showed a a strong genotype effect in both females and males. Data presented as the mean binding intensity +/- SEM, with data points for each individual mouse included. **p*-value = 0.05, ***p*-value = 0.01, *****p*-value = 0.0001.

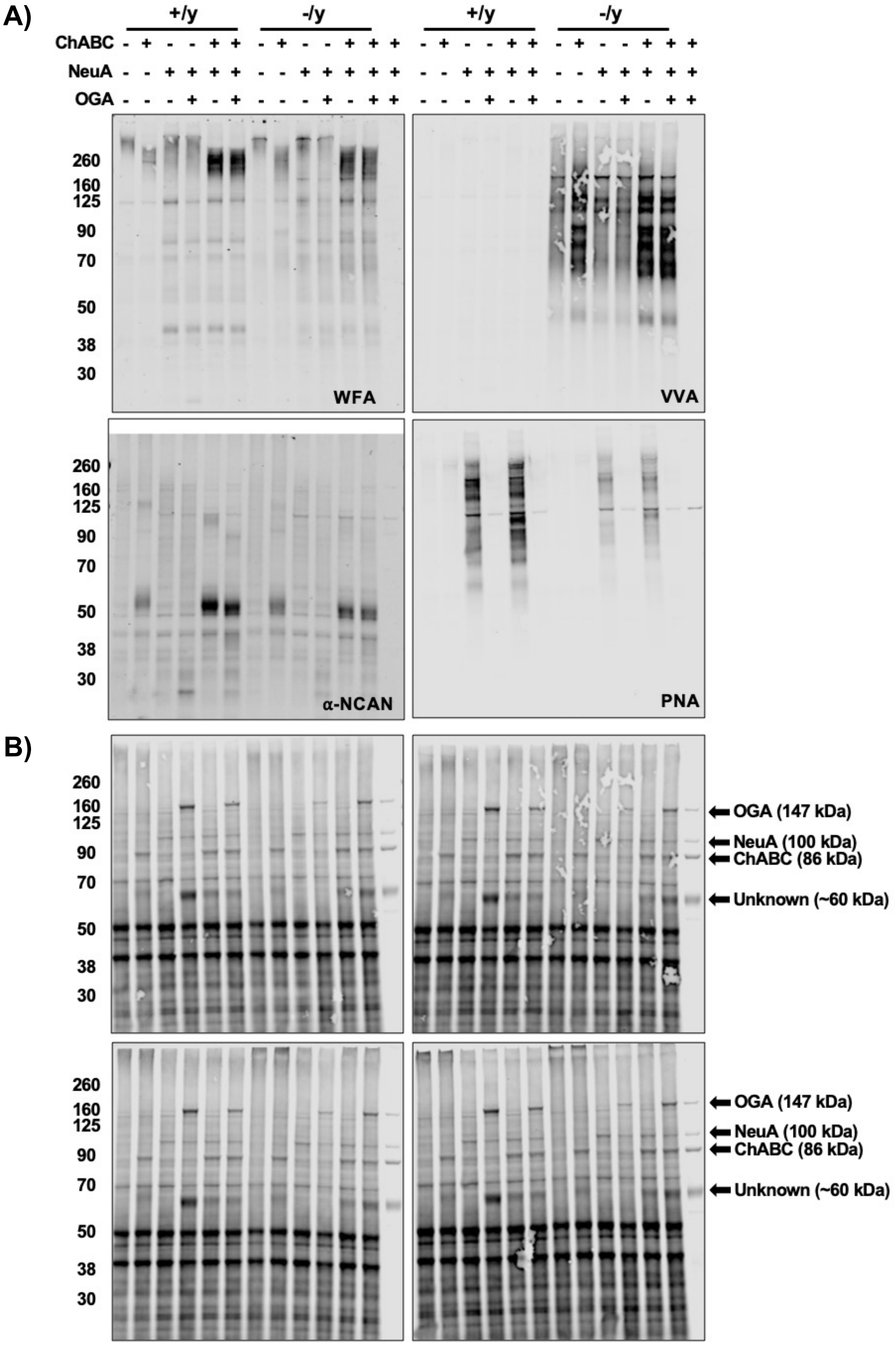


**Supplemental Figure 4. O-GalNAc and chondroitin sulfate modifications are present on the same protein carriers.** Lectin and antibody blotting following combinatorial glycosidase treatments in WT (+/y) and *Slc35a2*-deleted (-/y) mouse cortex. The migration pattern of WFA+ bands shows smaller molecular weight upon treatment with chondroitinase ABC (ChABC) in WT cortex. Minimal effect is seen with NeuA alone. However, treatment with NeuA and ChABC results in a dramatic increase in WFA signal, and no major change was seen with OGA treatment. In -/y cortex, the pattern is similar while the overall intensity of WFA binding is reduced. Blotting for NCAN, a classic CSPG that also carriers O-GalNAc glycans shows a similar pattern of increasing with ChABC + NeuA. However, addition of OGA reduces the apparent molecular weight of NCAN, consistent with both CPSG and O-GaNAc glycans being present on the same protein carrier. In -/y mice, the overall intensity of NCAN binding is reduced, but there is no apparent size difference following treatment with OGA, suggesting that O-GalNAc glycans are deficient on NCAN while proteoglycans remain relatively intact. VVA binding increases with ChABC treatment in -/y mice, and even more so upon treatment with NeuA, further confirming the complex interplay of these modifications on the same protein. VVA binding is insensitive to all glycosidases tested. Finally, PNA binding confirms NeuA activity and the reduction in -/y cortex. B) Total protein loads and bands corresponding with glycosidase input signals, including OGA (147 kDa), NeuA (100 kDa), ChABC (86 kDa subunit). Another prominent band at 60kDa is present in the enzyme loading control of unknown origin. Each lane contains 15 µg cortical protein lysate from an individual mouse of the corresponding genotype.

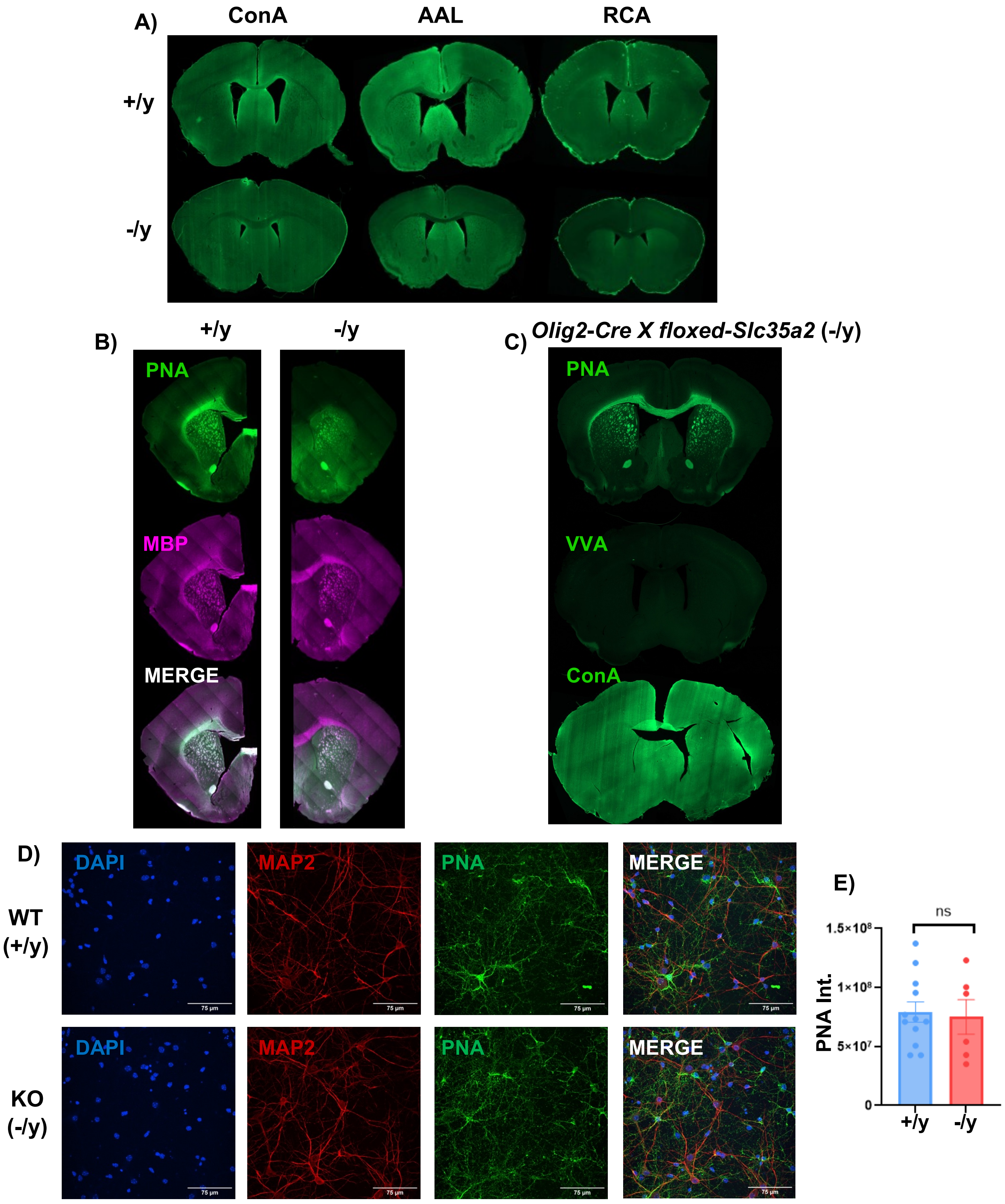


**Supplemental Figure 5. Cortical *Slc35a2* deletion does not impact the distribution of N-glycans or white matter tracts of the corpus callosum. A)** Coronal mouse brain sections of ConA, AAL, and RCA binding in +/y and -/y mice shows minimal difference between genotypes. **B)** Co-labeling with PNA and an antibody for myelin basic protein (MBP) shows near complete overlap in white matter tracts of +/y mice, while -/y mice lack PNA binding in the CC even though MBP labeling remains intact. **C)** *Olig2-Cre X floxed-Slc35a2* (-/y) mouse brains showed normal PNA and ConA binding and were VVA negative. **D)** Loss of *Slc35a2* in primary mouse neurons does not change total O-GalNAc levels. PNA fluorescent images of +/y and -/y primary neuron cultures at DIV11. Scale bar = 75 µm. **B\E)** Quantification of raw integrated density values of overall PNA fluorescence. ROIs were selected via automated module. N = 3 independent cultures/genotype. Scale bar = 10 µm.

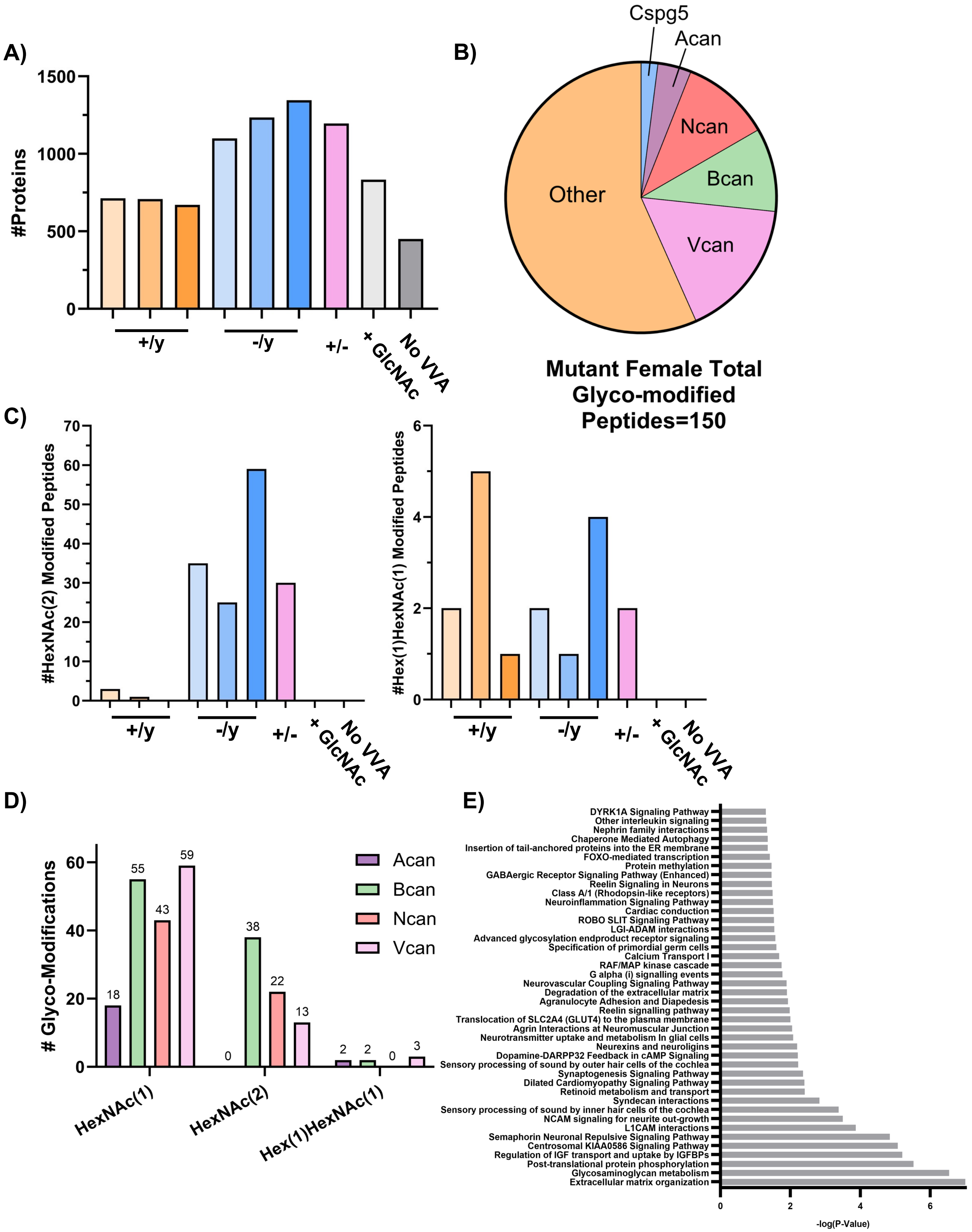


**Supplemental Figure 6. Extended glycoproteomic data of *Slc35a2*-deleted cortex. A)** The total number of proteins with spectral count > 0 detected in each sample, highlighting the presence of non-specific binders. **B)** HexNAc modified glycopeptides from male (n = 3) and female (n = 1) enrich on lecticans in both sexes. **C)** HexNAc(2) modified peptides (+406.158 Da) were detected almost exclusively in the *Slc35a2*-deleted cortex, and very few Hex(1)HexNAc(1) modified peptides (+365.132 Da) were detected across all samples. **D)** All lecticans had a majority of HexNAc(1) modifications. **E)** Gene Ontology (GO) analysis of the 60 mapped genes that contained the 443 detected glycopeptides revealed several significantly enriched pathways, most notably GAG metabolism and the ECM.

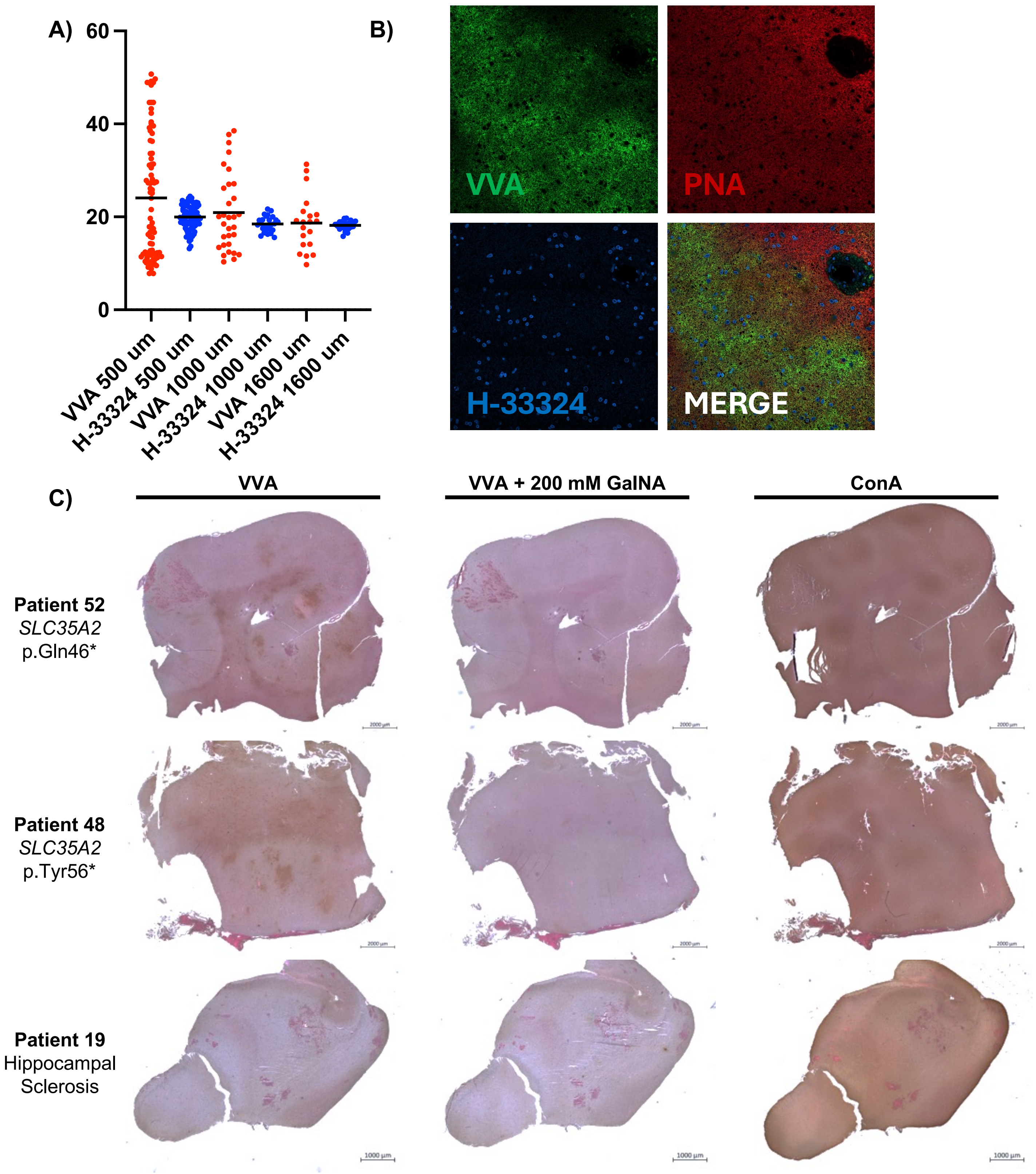


**Supplementary Figure 7. Additional VVA staining in human DRNE cases. A)** Sub-millimeter variability in O-GalNAc density may impact tests of VAF in *SLC35A2*-associated epilepsy. Random 500 x 500 µm, 1,000 x 1,000 µm, and 1,600 x 1,600 µm fluorescent intensity measures of VVA and H-33324 signal from Fig. 5E, plotted as arbitrary intensity (a.i.) normalized to average background signal adjacent to tissue block. Bar indicates median, N = 82, 80, 31, 26, 20, and 23, respectively. **B)** VVA and PNA binding appear most intense in separate locations of a high mutation case (VAF = 62.6%). Scale bar = 25 µm. **C)** Histochemical identification of Tn Antigen human tissue. Representative images of tissue samples stained with VVA and visualized using DAB histochemistry. Patient tissue harboring pathogenic loss of function variant in *SLC35A2* displays significant VVA positivity, while control tissue from an individual with a non-*SLC35A2*-associated epilepsy (Hippocampal Sclerosis, *SLC35A2* VAF = 0%) does not. Lectin specificity is confirmed by competitive inhibition of VVA binding using 200 mM GalNAc treatment. Major changes to N-glycans detected by ConA are not apparent, complementary to our other results. Scale bar (Patient 52 & 48) = 2000 µm. Scale bar (Patient 19) = 1000 µm.

A)

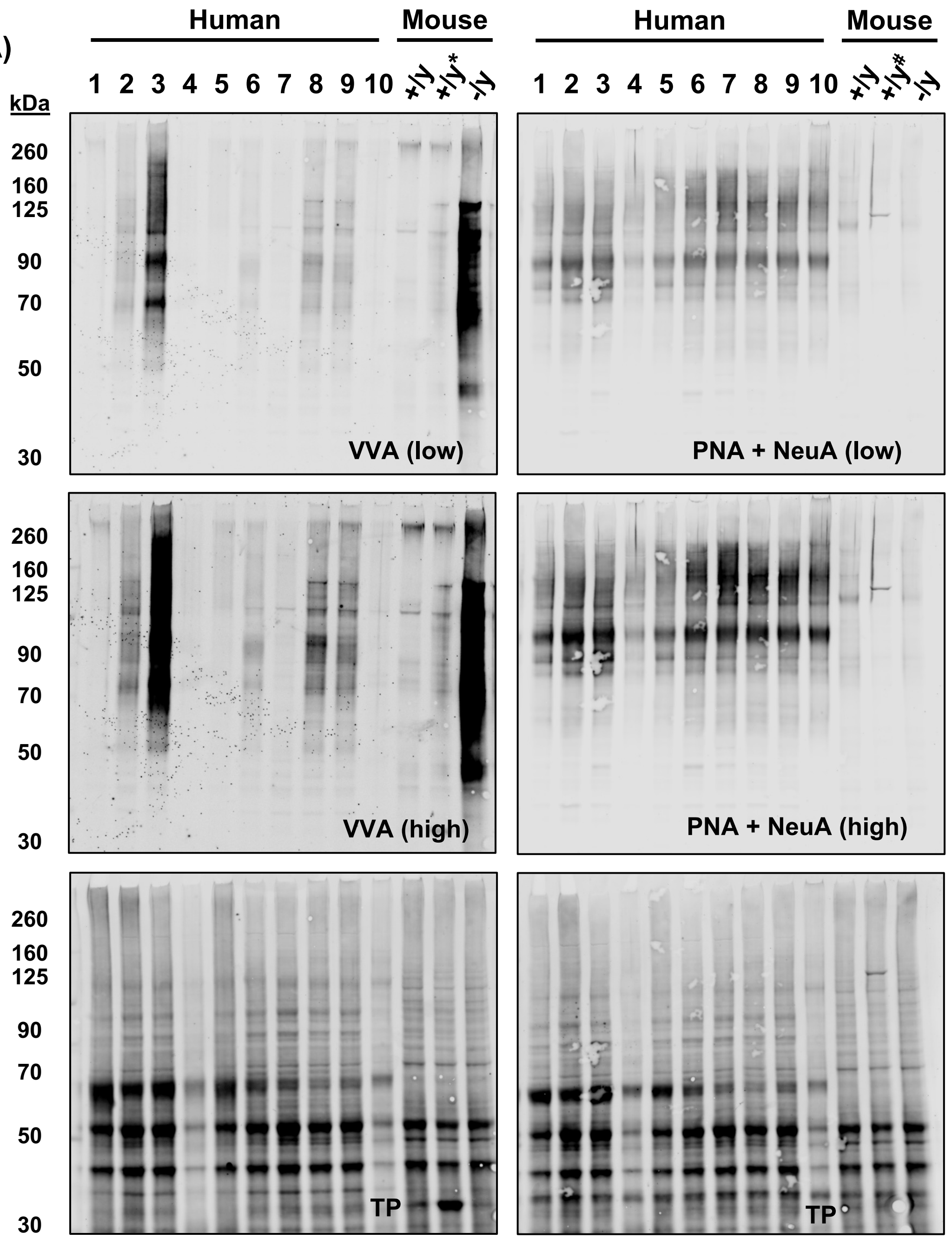


B)

| Sample | 1 | 2 | 3 | 4 | 5 | 6 | 7 | 8 | 9 | 10 | | |
|---|---|---|---|---|---|---|---|---|---|---|---|---|
| Disorder | MTOR-NPRL3 | SLC35A2 | SLC35A2 | MTOR-DEPDC5 | SLC35A2 | SLC35A2 | MTOR-MTORA1 | SLC35A2 | SLC35A2 | MTOR-MTORA1 | | |
| VVA | - | + | + | - | - | + | - | + | + | - | **r^2** | ***p*-value** |
| *Reported VAF % | 0.0 | 3.0 | 27.7 | 0.0 | 2.5 | 11.0 | 0.0 | 6.3 | 21.4 | 0.0 | **0.669** | **0.0131** |
| Measured VAF % | 0.0 | 0.5 | 16.8 | 0.0 | 0.2 | 2.5 | 0.0 | 5.2 | 3.4 | 0.0 | **0.973** | **<0.0001** |

**Supplemental Figure 8. Blind analysis of human epilepsy brain tissue using VVA correctly identified 80% of *SLC35A2*-associated epilepsy in cohort 1. A)** Human brain lysate from tissue removed during brain biopsy for intractable epilepsy correctly identified 80%(4/5) of cases, only missing the sample with the lowest known variant allele frequency (VAF) of 2.5%. No other epilepsy cases were determined to be VVA positive (0/5, 0%). Cortical lysate of *Slc35a2*-Emx1 mice were included as VVA negative (+/y), VVA positive (-/y), and enzyme controls (*PNGase F, #OGA)(right 3 lanes). Low (top row) and high (middle row) exposure images for VVA and PNA + NeuA are shown, as well as the corresponding total protein (TP, bottom row) load. Cortical lysate of *Slc35a2*-Emx1 mice were included as negative (+/y) and positive (-/y) VVA controls (right 3 lanes). 15 µg of brain protein lysate was loaded per lane. **B)** Results from A summarized with the corresponding disorder type, initial VAF% calculation, and whether the same was deemed VVA positive (+) or negative (-) from a blind reviewer. *Initial VAF% calculation was performed on an adjacent but separate tissue block.

A)

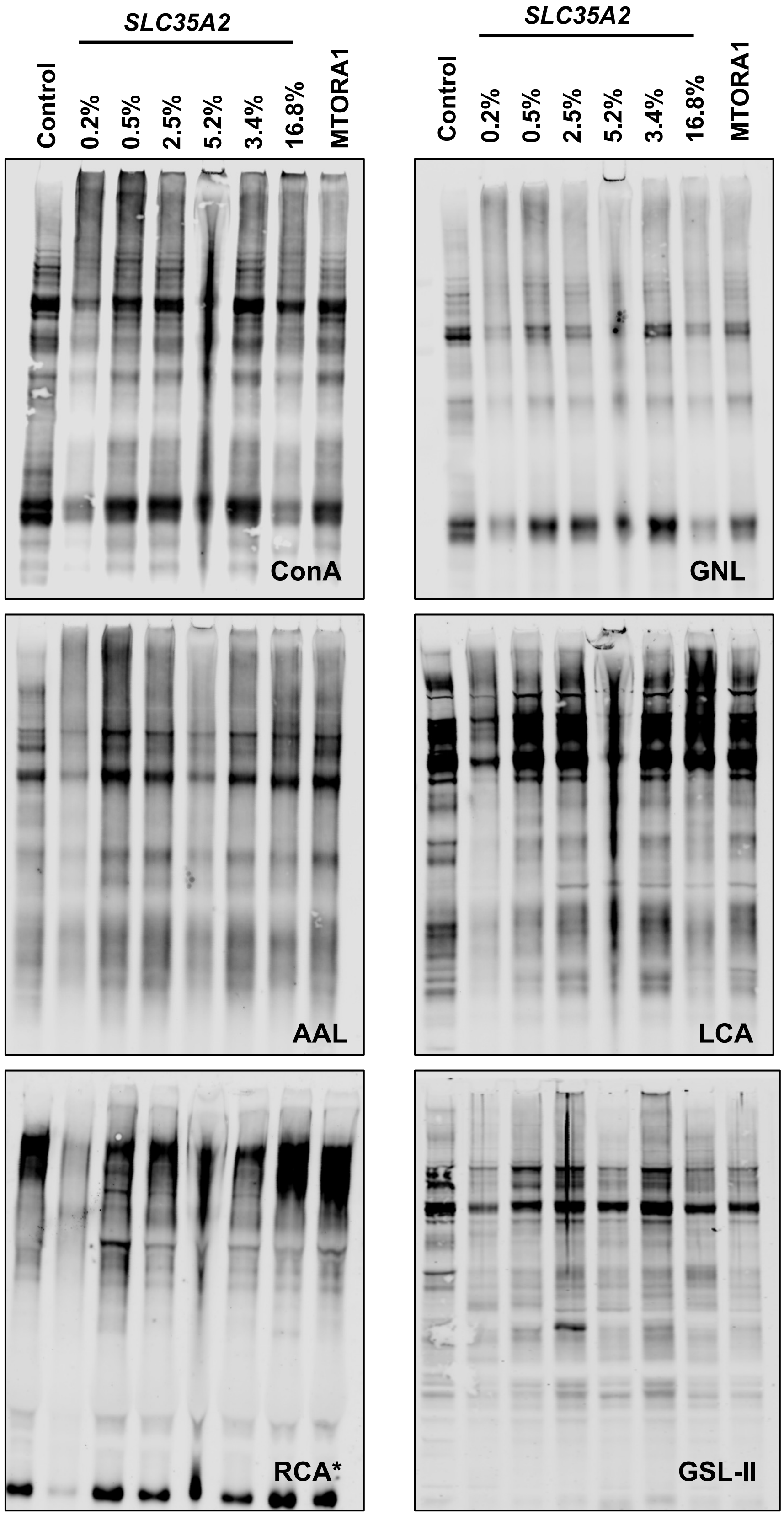


B)

| Lectin | VVA/TP | PNA* | ConA | GNL | AAL | LCA | RCA* | GSL-II |
|---|---|---|---|---|---|---|---|---|
| **r2** | 0.973 | 0.006 | 0.263 | 0.193 | 0.008 | 0.001 | 0.129 | 0.072 |
| **p-value** | **<0.0001** | 0.852 | 0.194 | 0.277 | 0.832 | 0.933 | 0.381 | 0.521 |

**Supplementary Figure 9. N-glycan binding lectins do not correlate with variant allele frequency (VAF) in acquired *SLC35A2* epilepsy. A)** Blots of N-glycan-binding lectins to human brain lysate from tissue removed during brain biopsy. *RCA was tested in the presence of α1-3,4-fucosidase to expose any potential galactose binding motifs. 15 µg of brain protein lysate was loaded per lane, with quantification of lectin binding normalized to total protein load. **B**) While VVA binding correlates with variant allele frequency (VAF) in acquired SLC35A2 epilepsy (Fig. S16) no other lectins tested showed a significant correlation to VAF%.

A)

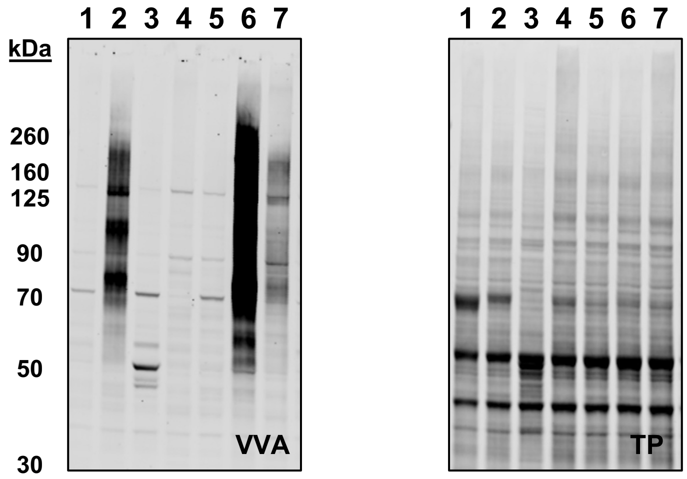


B)

| Cohort 2 | 1 | 2 | 3 | 4 | 5 | 6 | 7 | | |
|---|---|---|---|---|---|---|---|---|---|
| Disorder | DRNE | SLC35A2 | DRNE | DRNE | DRNE | SLC35A2 | SLC35A2 | | |
| VAF % | 0 | 19.7 | 0 | 0 | 0 | 62.6 | 3.0 | **r^2** | ***p*-value** |
| VVA | - | + | - | - | - | + | + | **0.99** | **<0.0001** |

**Supplemental Figure 10. Blind analysis of a second cohort of *SLC35A2*-associated correctly identified 100% of cases based on VVA positivity . A)** Human brain lysate from tissue removed during brain biopsy for intractable epilepsy correctly identified 100% (3/3) of *SLC35A2* cases. No other epilepsy cases, which are histologically confirmed as FCD 2b but lack a genetic diagnosis, were VVA positive (0/4, 0%). Total protein (TP), right panel. 15 µg of brain protein lysate was loaded per lane. **B)** Results from A summarized with the corresponding disorder type, initial VAF% calculation, and whether the sample was deemed VVA positive (+) or negative (-) from a blind reviewer. *Initial VAF% calculation was performed on an adjacent but separate tissue block.
